# A haplotype-resolved, chromosome-scale genome assembly for the southern live oak, *Quercus virginiana*

**DOI:** 10.1101/2025.08.26.670957

**Authors:** Laramie Aközbek, Zachary Meharg, Jillian Abendroth-McGhee, Tosin Akinsipe, Rijan Dhakal, Nicholas Gladstone, Zahida Pervaiz, Sejal Patel, Giovani Rossi, Claudia Ann Rutland, Caroline Bendickson, Adam Kranz, Ellen O. Martinson, Scott P. Egan, F. Alex Feltus, David J. Clarke, Jeremy Schmutz, John Lovell, Jenell Webber, Lori Beth Boston, Haley Hale, Hannah McCoy, Jane Grimwood, Sarah B. Carey, Leslie Goertzen, Alex Harkess

## Abstract

Hybridization is a major force driving diversification, migration, and adaptation in *Quercus* species. While population genetics and phylogenetics have traditionally been used for studying these processes, advances in sequencing technology now enable us to incorporate comparative and pan-genomic approaches as well. Here we present a highly contiguous, chromosome-scale and haplotype-resolved genome assembly for the southern live oak, *Quercus virginiana*, the first reference genome for section *Virentes*, as part of the American Campus Tree Genomes (ACTG) program. Originating from a clone of Auburn University’s historic “Toomer’s Oak,” this assembly contributes to the pool of genomic resources for investigating recombination, haplotype variation, and structural genomic changes influencing hybridization potential in this clade and across *Quercus*. It also provides insights into the architecture of the putative centromeric regions within the genus. Alongside other oak references, the *Q. virginiana* genome will support research into the evolution and adaptation of the *Quercus* genus.

## Introduction

Hybridization has long been recognized as a powerful evolutionary force in oaks (*Quercus* spp.) that has fueled their diversification and migration across the globe as well as their adaptation to new environments (Kremer and Hipp 2020). A high potential for interfertility allows oaks to form syngameons, which are systems where three or more interbreeding species living in sympatry are able to retain their distinctiveness despite repeated interspecific hybridization and introgression. Although suspected to occur in other sections of *Quercus*, the syngameons of subg. *Quercus* sec. *Quercus* (∼150 species) are the most well-studied. The Southern Live Oak (*Quercus virginiana* Mill.) is a member of subg. *Quercus* sec. *Virentes* (7 species), which is sister to sec. *Quercus*, and is narrowly distributed in both North America and Cuba (Cavender-Bares et al. 2015). Members of this small section have a strong history of within-section hybridization, but are not frequently sympatric when compared to more species-rich sections. As a result, the species in sec. *Virentes* are generally not associated with a syngameon, but their participation in one still remains an open question (Cavender-Bares et al. 2015; Eaton et al. 2015; Cannon et al. 2024). However, the existence of natural and artificial hybrids between sec. *Quercus* and sec. *Virentes* (i.e. Compton’s Oak, *Quercus* x *harbsonii*, etc.) indicate that reproductive barriers are minimal between them (International Oak Society, Date Unknown; Nesom 2018).

The propensity for hybridization across the genus has driven researchers to utilize population genetics and phylogenetics to examine how gene flow influences the population structure of oaks and other species, shapes their evolutionary history, and enhances their adaptive potential (Eaton et al. 2015; McVay et al. 2017; Cavender-Bares 2019; Whittemore and Miller 2023). However, a comprehensive characterization of the oak syngameon requires the synthesis of population genetic and genomic approaches (Cannon and Petit 2020). High-quality reference genomes and pan-genomes are essential to understanding this phenomenon, allowing for a more accurate and precise description of genetic variation in target species. As genomic architecture plays a critical role in the maintenance of interfertility and extensive chromosomal rearrangements between species can impact hybridization potential, comparative genomics can help reveal the genetic mechanisms underlying reproductive barriers, the extent of structural variation between and within species, the impact of this variation on gene flow, and the evolutionary dynamics shaping species divergence (Rieseberg 2001).

To help address these broad questions, we present a chromosome-scale, haplotype-resolved diploid assembly for *Quercus virginiana*, a species of cultural and ecological importance in the southeastern United States. This genome was generated as part of the American Campus Tree Genomes (ACTG) program (www.hudsonalpha.org/actg), where undergraduate and graduate students assemble, annotate, and publish tree genomes from their college campuses. This *Q. virginiana* accession is a propagated clone of “Toomer’s Oak,” an important symbol of Auburn University that was tragically poisoned by the herbicide Spike 80DF (Tebuthiuron) in 2010. Phylogenetically, this assembly is the first representative genome of the oak section *Virentes*. As such, it contributes to the growing genomic resources for *Quercus*. Reference genomes like the one we report here support and enrich population genetic research, as the variation between haplotypes allows researchers to identify hundreds, often thousands, of loci with distinct histories. Haplotype-resolved genomes also allow for rigorous investigation into macro and micro-structural variation between haplotypes within and between species. Since structural changes, such as inversions, translocations, and deletions, are suspected to play key roles in speciation, understanding these elements of the *Q. virginiana* genome will give insights into future investigations of genome evolution and species boundaries in sec. *Virentes* (Berdan et al. 2024). As more high-quality assemblies become available for highly heterozygous, sympatric *Quercus* species, they will further contribute to our understanding of the genetic basis of hybridization dynamics and the syngameons of this genus.

## Methods

### Sample collection, extraction, and sequencing

To generate whole genome sequencing data to assess the heterozygosity and genome size of *Q. virginiana*, a standard CTAB method was used to isolate DNA from young leaf tissue. 3 micrograms of input DNA was used to construct Illumina TruSeq DNA PCR-free libraries and these libraries were subsequently sequenced on an Illumina NovaSeq6000 using PE150 reads.

Approximately 20 grams of young leaf tissue was collected from a Toomer’s Oak clone and flash-frozen in liquid nitrogen for use in PacBio HiFi sequencing. A voucher is available at the Auburn University John D. Freeman Herbarium (AUA:71025). Using the Circulomics Nanobind Plant Nuclei Big DNA kit, high molecular weight DNA was isolated from young leaf tissue with 4 grams of input tissue and a 2-hour lysis. The high molecular weight DNA quality was assessed via spectrophotometry for purity, via the Qubit dsDNA Broad Range assay for concentration, and run on the Agilent Femto Pulse to check fragment size. A Diagenode Megaruptor was used to shear the DNA, which was then size-selected to roughly 25 kb on a BluePippin. The SMRTbell Express Template Prep Kit 2.0 was used to build the PacBio libraries with CCS (HiFi) reads generated using two PacBio Sequel-II 8M flow cells at the HudsonAlpha Genome Sequencing Center. A Dovetail Omni-C library was generated using 1g of flash-frozen tissue as input, following the manufacturer’s protocol, and sequenced on an Illumina NovaSeq6000 using PE150 reads. RNA was isolated from four vegetative tissue types (young leaves, brown and green senescing leaves and roots) using a modified CTAB approach followed by clean-up with a Zymo RNA Clean & Concentrator kit. RNA-seq libraries were generated using the Illumina TruSeq stranded mRNA kit following the manufacturer’s protocol, and sequenced on an Illumina NovaSeq6000 using PE150 reads.

### Genome assembly and scaffolding

HiFi read quality and distribution were assessed with *nanoplot* (v1.42.0) (De Coster et al. 2018). Ploidy and heterozygosity were estimated utilizing GenomeScope2 (v1.0.0) and Smudgeplot (v0.2.5) (Ranallo-Benavidez et al. 2020). The raw HiFi reads were assembled into contigs using *hifiasm* (v0.20.0-r639) with Omni-C integration (Cheng et al. 2021). The resulting haplotypes were then combined and polished with the HiFi long-reads using *racon* (v1.5.0) (Vaser et al. 2017). The assemblies were subsequently screened for contaminants with FCS-GX (v0.5.4), then *bbmap* (v39.13) was used to drop any contigs less than 25kbp from the assembly (Bushnell 2014 Mar 17; Astashyn et al. 2024). Prior to scaffolding the genome of *Q. virginiana*, we mapped the Omni-C reads to the preliminary assembly with *bwa mem* (v0.7.17, flags: -5SP -T0), then filtered for duplicate and unmapped reads with *pairtools* (v0.3.0) using default parameters (Li and Durbin 2009; Open2C et al. 2024). We scaffolded the assembly with *Yet another Hi-C Scaffolding Tool* (*YaHS*, v1.1) using default parameters (Zhou et al. 2023). We visualized and manually curated the assembly with Juicebox (v2.13.07) (Durand et al. 2016, Supplementary Figure S1). Telomeres were identified with *GENESPACE* (v1.3.1) to check for their presence or absence as well as to assess their proper orientation within the chromosome (Lovell et al. 2022). Genome completeness and quality metrics were assessed utilizing *assemblathon2, merqury* (v1.3), and *compleasm* (v0.2.6) with the lineage *eudicots_odb10* (Bradnam et al. 2013; Rhie et al. 2020; Huang and Li 2023). The *Q. virginiana* assemblies were reordered and named according to the previously published *Quercus* references (Kapoor et al. 2023). The plastid genomes were assembled from the raw HiFi reads with *OatK* (v1.0), annotated with *GeSeq* (v2.0.3), and visualized with *OGDRAW* (v1.3.1) (Tillich et al. 2017; Greiner et al. 2019; Zhou et al. 2024).

### Genome annotations

We ran *RepeatModeler2* (v2.0.6) with the *LTRStruct* parameter to generate a *de novo* repeat library for each haplotype, which was used to softmask the assembly with *RepeatMasker* (v4.1.5) (Chen 2004; Flynn et al. 2020). Repetitive elements were further annotated with *EDTA* using default parameters (v2.1.3) to examine the repeat landscape (Ou et al. 2019, Supplementary Figure S2). Centromeric monomers were identified with *TRASH* (v1) and centromeric arrays were visualized with *StainedGlass* (v0.6), *RepeatObserverV1*, and *karyoploteR* (v1.28.0) (Benson 1999; Gel and Serra 2017; Vollger et al. 2022; Elphinstone et al. 2023; Wlodzimierz et al. 2023). Additional information regarding the identification, characterization and visualization of the *Q. virginiana* centromeres can be found in Supplemental Methods S1 in the Supplementary Material.

We performed a gene annotation with *Braker3* (v3.0.6), utilizing proteins from other *Quercus* and Fagales species as well as RNA-seq evidence from various *Q. virginiana* tissue types: young leaves, brown and green senescing leaves and roots (Gabriel et al. 2023, Supplementary Table S1). Annotations without full or partial support from RNA-seq evidence were removed from the final gene set. The functional annotation was performed with *EnTAP* (v2.1.0) and the completeness of the annotation was assessed with *BUSCO* (v5.7.0) using the *eudicots_odb10* database (Hart et al. 2020; Manni et al. 2021).

### Structural and comparative genome analyses

The haplotypes were compared to each other with *nucmer* (v4.0.0rc1) and visualized with dot (Sandbox Bio, Date Unknown; Supplementary Figure S3). Structural variants between the haplotypes and other genome assemblies were called with *syri* (v1.7.0) and *plotsr* (v1.1.0) (Goel et al. 2019; Goel and Schneeberger 2022). *GENESPACE* (v.1.3.0) was used to build riparian plots to visualize syntenic relationships between the haplotypes as well as other *Quercus* species (Lovell et al. 2022).

## Results and Discussion

We present a chromosome-scale assembly for the diploid *Quercus virginiana* “Toomer’s Oak” using a combination of PacBio HiFi reads and Dovetail Omni-C (Supplementary Figure S4). Nearly 200x coverage of paired-end 150bp Illumina sequencing data was generated, and k-mer frequencies were used to estimate a haploid genome size of 719 Mb and 1.57% heterozygosity (Supplementary Figures S5 and S6, Supplementary Table S2). Approximately 66 Gb of raw PacBio HiFi data were generated, amounting to ∼84x estimated coverage of the haploid nuclear genome size estimate (Supplementary Table S2, Supplementary Figure S7). We assessed the quality of our HiFi reads using *nanoplot*, indicating high-quality libraries and a mean read length distribution centered around 14.5 kb (Supplementary Figure S8, Supplementary Tables S2 and S3). Of the ∼792M Omni-C read pairs generated scaffolding, there was a low duplication rate (Supplementary Table S4). Haplotype 1 was 788.8 Mb in length with a contig N50 of 53 Mb, and Haplotype 2 was 781.7 Mb in length with a contig N50 of 55.4Mb (Table 1, Supplementary Table S2). In both haplotypes, ∼50% of chromosomes are contained in single contigs, and all chromosomes are flanked with canonical telomeric sequences (Table 1, Figure 1b, Supplementary Figure S9). Assembly BUSCO scores are strong for both haplotypes, with ∼99% complete genes recovered (Table 1, Supplementary Table S2). *k*-mer-based completeness statistics indicate a high consensus quality score (QV > 43) and k-mer completeness score (98.1%) for the combined haplotypes (Table 1, Supplementary Table S2).

**Table 1.** Genome Assembly and Annotation Statistics

| Assembly |  |  |
| --- | --- | --- |
| Assembly Statistics | Haplotype 1 | Haplotype 2 |
| Total Contig Length (Mb) | 788.8 | 781.7 |
| Number of Contigs | 601 | 323 |
| Longest Contig (Mb) | 100.62 | 97.96 |
| Min. Number of Contigs containing half of assembly, L50 | 6 | 6 |
| Shortest Contig from L50 set, N50 (Mb) | 53.03 | 55.37 |
| Number of Scaffolds | 586 | 309 |
| Min. Number of Scaffolds containing half of assembly, L50 | 5 | 5 |
| Shortest Scaffold from L50 set, N50 (Mb) | 65.99 | 66.33 |
| Base pair QV | 43.754 | 43.9434 |
|  | both = 43.8473 |  |
| k-mer completeness (%) | 79.34 | 79.71 |
|  | both = 98.08 |  |
| Assembly BUSCO Completeness (%) |  |  |
| Complete (S + D) | 99.78 | 99.78 |
| Single-copy (S) | 96.73 | 96.9 |
| Duplicated (D) | 3.05 | 2.88 |
| Fragmented (F) | 0.09 | 0.09 |
| Missing (M) | 0.13 | 0.13 |
| Repeats (RepeatMasker) |  |  |
| LAI (LTR Assembly Index) | 20.61 | 21.16 |
| Repeats (%) | 61.21 | 61.17 |
| Annotation |  |  |
| Annotation Statistics | Haplotype 1 | Haplotype 2 |
| Number of Protein-coding Genes | 30,846 | 31,361 |
| Number of Protein-coding Transcripts | 36,129 | 36,559 |
| Annotation BUSCO Completeness (%) |  |  |
| Complete (S + D) | 97.6 | 97.2 |
| Single-copy (S) | 94.2 | 93.8 |
| Duplicated (D) | 3.4 | 3.4 |
| Fragmented (F) | 0.5 | 0.5 |
| Missing (M) | 1.9 | 2.3 |

**Figure 1.**
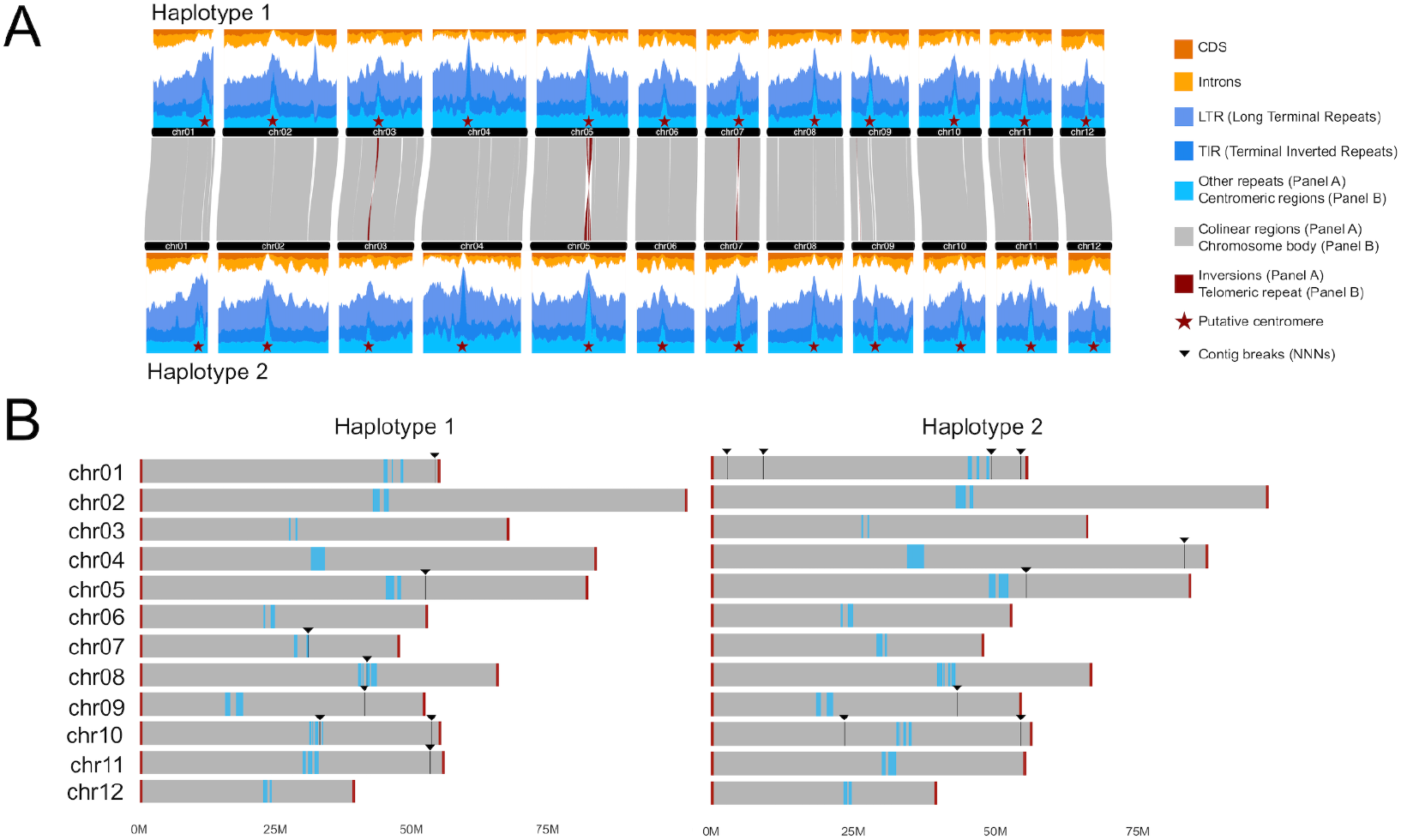
The genomic structure of *Quercus virginiana*. **(A)** the gene and repeat landscapes of both haplotypes. Although the trend is not as pronounced in other angiosperms, repeat density tends to increase and gene density tends to decrease as both approach the centromeres. **(B)** a graphical representation of the *Q. virginiana* karyotype for both haplotypes, showing the proximity of breaks in contiguity to the putative centromeric satellites as well as the presence of canonical telomeric sequences flanking all chromosomes.

The final assembly is haplotype-resolved with 12 chromosomes per haplotype. Structural variation between the haplotypes, as characterized by *syri*, consists primarily of 54 inversions, approximately 244 indels and 3.29M SNPs, as well as a varying number of small duplications and translocations (Supplementary Table S5). We did not find any known contaminants, and furthermore, we searched for evidence of horizontal gene transfer (HGT) from the cynipid gall wasp *Belonocnema kinseyi*, a gall-inducing wasp hosted by *Q. virginiana*, as well as bacterial and fungal sources into the *Q. virginiana* genome, but did not find any evidence for HGT between the oak and these symbionts (Supplementary Methods S2). We also assembled and annotated the chloroplast genome of *Q. virginiana* (Supplementary Figure S10). The size of the chloroplast genome (161kb) is similar to that of previously published assemblies with 80 protein-coding, 30 tRNA and 4 rRNA annotations (Yang et al. 2016; Hu et al. 2019; Zhang et al. 2020).

After filtering, Haplotype 1 contains 30,846 predicted protein-coding genes, while Haplotype 2 contains 31,361 with BUSCO completeness scores of 97.6% and 97.2%, respectively. Approximately 81% of protein-coding genes are functionally annotated (Supplementary Table S2). The repetitive content of both haplotypes is comparable, with approximately 61% of each haplotype annotated as repetitive (Table 1, Supplementary Table S2). Retrotransposons account for 24.89% of Haplotype 1 and 24.93% of Haplotype 2 while DNA transposons represent 19.84% and 20.8% of each haplotype, respectively (Supplementary Table S2). As seen in many other angiosperm genome assemblies, gene density decreases and repeat density increases as both approach the putative centromeres (Figure 1a). The major structural variants found between the haplotypes are inversions, particularly at the putative centromeric locations of Chr03, Chr05, Chr07, and Chr11 (Figure 1a).

### Comparative Genomics

Numerous reference genomes for diverse *Quercus* species have been published in recent years, presenting an exciting opportunity to explore the genomic basis of reproduction in the genus using comparative genomics. The genome of *Q. virginiana* is approximately 780Mb in size, which is close to the average genome size of other published *Quercus* assemblies: 818Mb (Plomion et al. 2018; Ai et al. 2022; Han et al. 2022; Sork et al. 2022; Zhou et al. 2022; Kapoor et al. 2023; Rey et al. 2023; Wang et al. 2023; Luo et al. 2024; Mead et al. 2024; Larson et al. 2025, Supplementary Table S6). Alongside *Quercus variabilis*, it represents one of the most contiguous and complete assemblies within the genus, exhibiting some of the highest assembly BUSCO completeness scores reported to date (Supplementary Table S6). While its repeat content is comparable to that of more recent assemblies, the total number of predicted protein-coding genes falls on the lower end of the range observed in other *Quercus* genomes, likely due to stringent filtering criteria applied during annotation and limited sampling of diverse tissues (Supplementary Table S6).

### Centromere positioning

Centromere position and structure may influence recombination dynamics in plants, potentially impacting adaptation, introgression, and other evolutionary processes in a clade (Garg et al. 2024). An analysis of the putative centromeres in *Q. virginiana* suggests that Chromosome 1 is acrocentric—a trait likely shared with other oak species such as *Quercus lobata* (Sork et al. 2022, Figures 1 and 2). The remaining chromosomes, however, exhibit varying degrees of metacentricity (Supplementary Figures S11 and S12). Satellite sequences associated with these centromeres display a “patchwork” distribution across certain chromosomes, flanking genic regions (Figure 2, Supplementary Figures S11 and S12, Supplementary Table S7). This pattern is unlikely to be the result of assembly artifacts, since only 3 of the 24 identified centromeres correspond with breaks in contiguity (Figure 1b) and similar forms appear in other highly contiguous *Quercus* genomes despite extensive structural rearrangements in the region (Figure 2b, Supplementary Figure S13). It is notable that over 11.3 Mb of sparsely genic sequence, containing approximately 240 genes, resides within these regions on Haplotype 1 of *Q. virginiana* (Supplementary Table S8).

**Figure 2.**
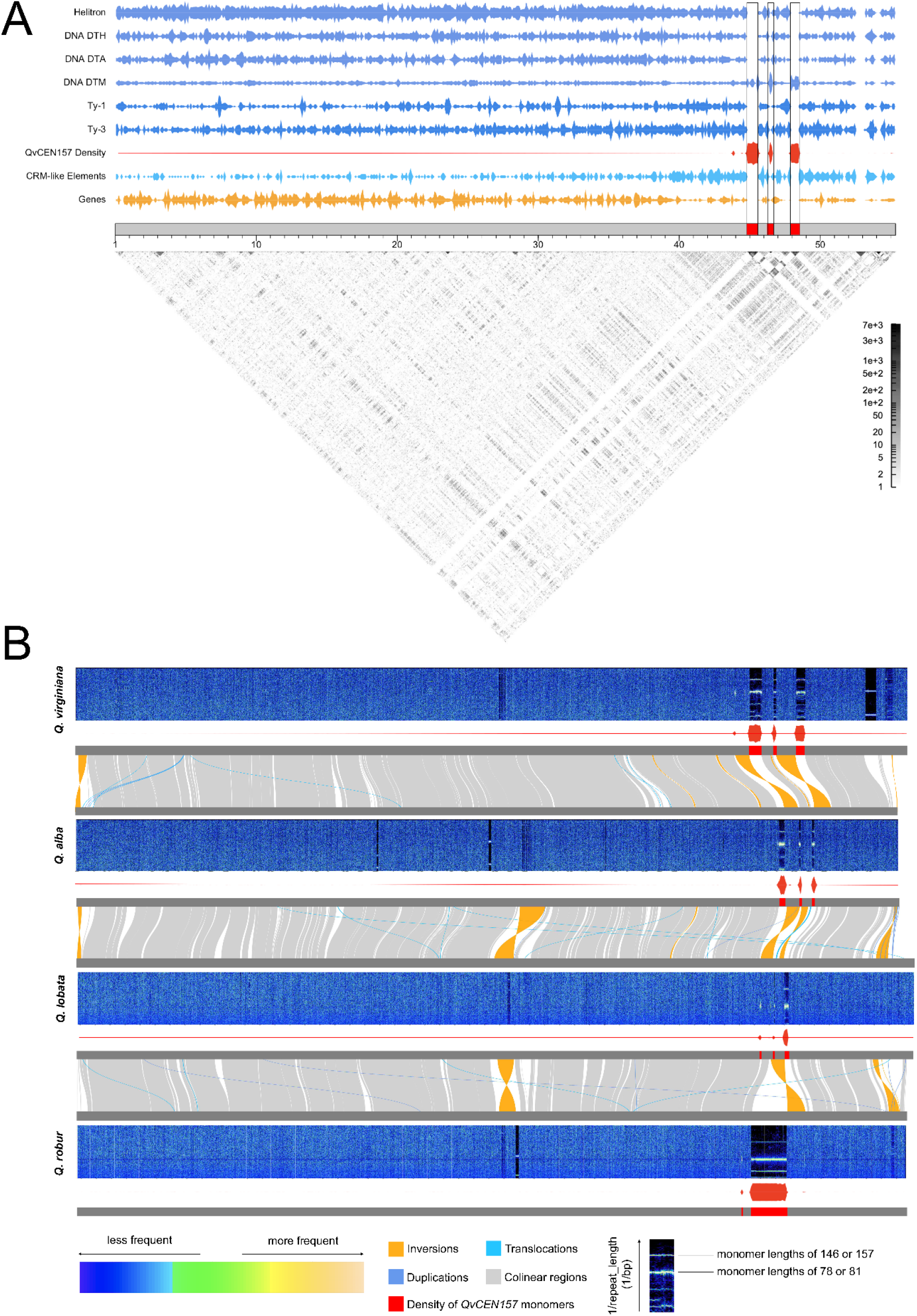
Putative Centromeric Satellites in Chromosome 1 of *Q. virginiana* and other North American and Eurasian sec. *Quercus* species. **(A)** the relative densities of diverse genomic features, including DNA and RNA transposable elements (blue), Helitrons (blue), genes (orange), and the *QvCEN157* monomer (red) with an identity heatmap (grey) generated with StainedGlass visualized in relation to the predicted centromeric satellites (red) (**B)** chromosome-scale Fourier spectra of Chromosome 1 across diverse *Quercus* species generated by RepeatObserver combined with a riparian plot showing base pair synteny between species as well as the density of *QvCEN157* monomers along each chromosome. The x-axis of the spectra indicates genomic position and the y-axis of the spectra represents the frequency of a particular repeat and the corresponding color intensity showing the abundance of the repeat at that frequency in that position of the genome. The uppermost bar is described as the true or “fundamental” frequency, and lower bars represent the harmonics of that true frequency. The top bar across these species either corresponds to 1/146 or 1/156:1/157, which agrees with the repeat monomers identified in *Q. virginiana* by TRASH. The presence of transposons can appear as “blurs” or “streaks” within and between the bars in the predicted array.

This patchwork arrangement on Chr1 is also observed in other North American section *Quercus* species, such as *Q. lobata* and the recently assembled *Q. alba*, but not in the Eurasian white oak, *Q. robur* (known as the “pedunculate oak” or “English oak), where a single satellite is present on all chromosomes (Wellcome Sanger Institute 2022, Figure 2b, Supplementary Figure S13). These putative centromeric satellites in *Q. virginiana* are associated with two closely related monomers: *QvCEN157* and *QvCEN146* (Supplementary Figure S14, Supplementary Tables S9 and S10). *QvCEN157* is associated most with chromosomes 1, 2, 3, 4, 5, 6, 7, 9, and 11, whereas *QvCEN146* is found mainly on chromosomes 8, 10, and 12 (Supplementary Figure S15). These monomer sizes are close to the reported putative centromeric monomer sizes in other oaks, such as *Q. lobata* (146 bp) and *Q. robur* (148 bp), although unsurprisingly, their sequence compositions differ (Sork et al. 2022;Xian et al. 2025, Supplementary Figure S16). Despite this, the *Q. virginiana QvCEN157* sequence can still be used to identify putative centromeric arrays in other species via a permissive *BLASTn* search (Figure 2b).

In *Q. virginiana*, a group of DNA transposons known as Mutator elements (DNA/DTM) have invaded the putative centromeric satellites to varying degrees, particularly in those of chromosome 4 (Figure 2a, Supplementary Figure S17). Centrophilic transposable elements have been documented in plants, however, they are typically retrotransposons (Naish and Henderson 2024). Centrophilic DNA transposons, which propagate without an RNA intermediate, are comparatively underrepresented in the literature. Mutator elements are well known for their mutagenic capabilities and are frequently linked to reproductive processes in plants (Ma and Li 2018; Dupeyron et al. 2019; Huang et al. 2024). This includes *Mutator-like element* (*MULE*) transposase-derived transcription factors such as *FAR1* (*FAR-RED IMPAIRED RESPONSE 1*) and *FSR (FAR1-RELATED SEQUENCE)*, the latter of which is found within the sparsely genic regions of the satellites on Chr1 in *Q. virginiana* (Supplementary Table S8).

Although the multi-satellite patchwork pattern is especially pronounced on chromosome 1 of this species and other taxa, the remaining chromosomes in the *Q. virginiana* genome also exhibit multiple putative centromeric satellites as do other *Quercus* lineages (Supplementary Figure S12). However, some of these other lineages, such as *Q. robur*, do not exhibit multiple satellites. The architecture described in *Q. virginiana* and other species is notable because it has not been well-described in the literature beyond a handful of recently assembled genomes, though it may be more widespread than currently recognized (Liu et al. 2023; Gao et al. 2024). Due to the presence of interspersed genic regions, this pattern does not conform to the classic monocentric model found in many angiosperms (Naish 2024). If CENH3 (Centromere-specific Histone H3) is associated with more than one of these satellites, suggesting di-centricity or tri-centricity, it is possible that proximity to each other enhances the stability of these regions, allowing them to behave similarly to a monocentromere, as in the stable di-centric rice centromere (Wang et al. 2013; Cuacos et al. 2015). Without additional experimental evidence, we also cannot classify these chromosomes as metapolycentric, a phenomenon observed in legumes such as *Pisum*, where multiple centromeric chromatin domains can be found across a primary constriction (Macas et al. 2023; Naish and Henderson 2024).

Recent research in *Arabidopsis* has revealed significant centromeric structural diversity both within and between species with some lineages undergoing substantial expansions of their centromeric satellites, yet CENH3 consistently localizes to only 1–2 Mb of centromeric regions, regardless of array’s size or composition (Naish 2024). This suggests that CENH3 localization is tightly regulated and reinforces the idea that centromere identity is epigenetically defined rather than dictated by genomic architecture. Given that CENH3 may localize to all, some, or none of these satellite sequences in *Q. virginiana* and other oaks, further experimental validation is necessary to characterize these regions. Continued advances in sequencing technology, particularly improvements in read length, accuracy, and assembly algorithms, will provide deeper insights into *Quercus* centromere evolution. As more genomes are assembled, we will have greater insights into the relationship between the evolution of these species and their centromeric architectures.

### Conserved gene order across Quercus

Additional comparative analyses between *Q. virginiana* and other *Quercus* species reveal a striking degree of synteny across divergent lineages, particularly between members of sec. *Quercus* and sec. *Virentes* (Figure 3). At this point in time, the only remaining clade lacking a representative genome is sec. *Ponticae*, but it is likely that it is also highly syntenic with the other sections. Larger structural variants are predominantly located near putative centromeric regions or at the distal ends of chromosome arms (Figure 3). The apparent synteny observed among *Quercus* species has been hypothesized to be both a cause and a consequence of the syngameon as conserved gene order may facilitate hybridization by reducing the likelihood of chromosomal incompatibilities, since extensive structural rearrangements are known to hinder gene flow between related taxa (Rieseberg 2001;Hipp et al. 2019; Cannon and Petit 2020; Rieseberg 2001). However, this observed conservation may not fully capture the extent of genomic diversity between or within species. To better understand the diversity of genomic architecture and its role in shaping the evolutionary dynamics within the genus, comprehensive pan-genomic approaches are needed.

**Figure 3.**
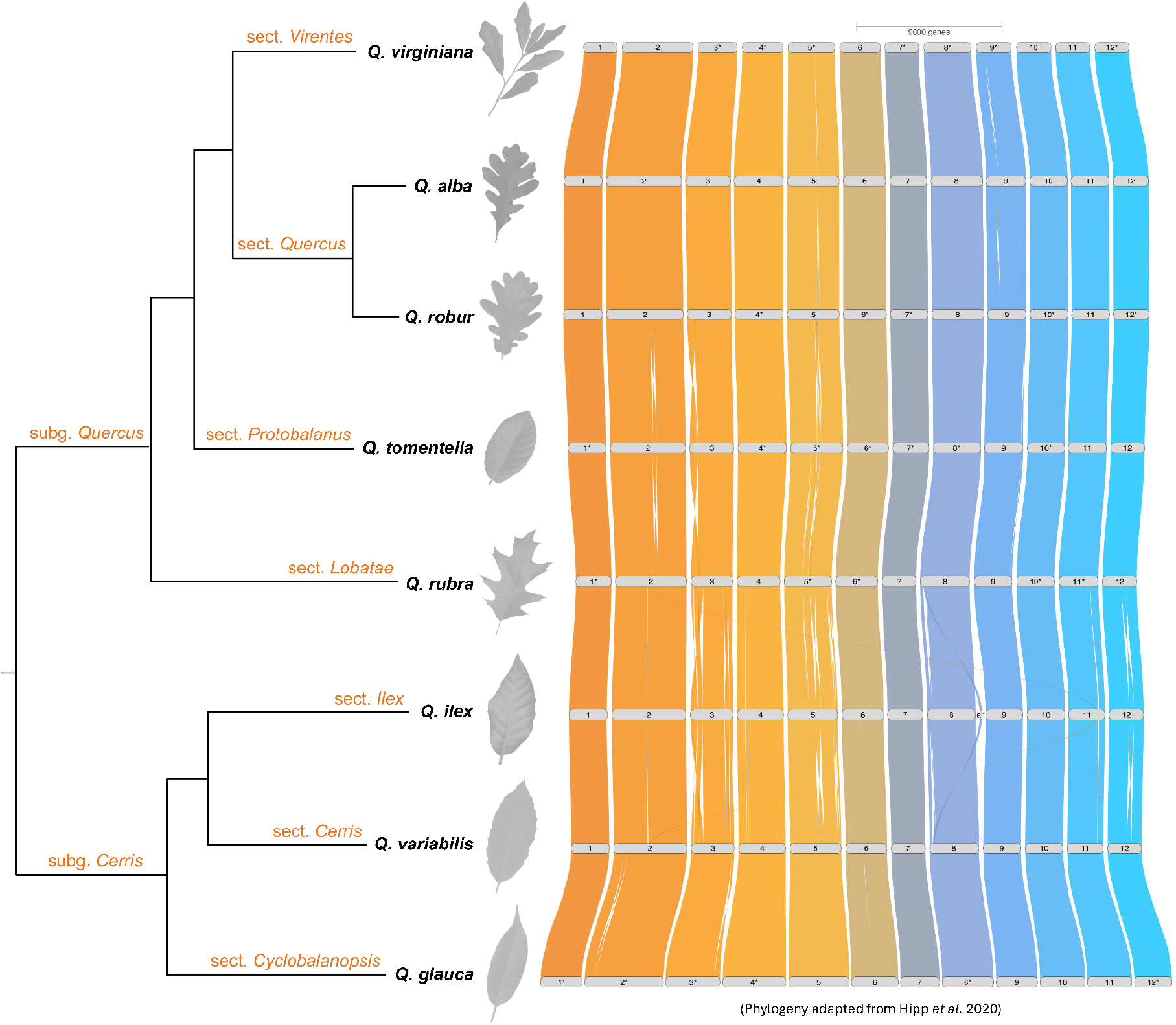
Riparian plot demonstrating conserved gene order between representative genomes from major sections of the genus *Quercus*, including *Q. alba* (Larson et al. 2025) and *Q. robur* for section *Quercus* (Wellcome Sanger Institute 2022), *Q. tomentella* (Mead et al. 2024) for section *Protobalanus, Q. rubra* (Kapoor et al. 2023) for section *Lobatae, Q. ilex* subsp. *ballota* (Rey et al. 2023) for section *Ilex, Q. variabilis* (Han et al. 2022) for section *Cerri*s, and *Q. glauca* (Luo et al. 2024) for section *Cyclobalanopsis*. Asterisks (*) indicate chromosomes that have been inverted relative to their original orientation for ease of visualization. The corresponding phylogeny was adapted from (Hipp et al. 2020).

## Conclusion

This chromosome-scale, phased diploid genome assembly of *Quercus virginiana* represents the first high-quality genomic resource for section *Virentes*. As one of the most contiguous and complete oak genomes available, it offers new insights into the structural organization of oak centromeres, revealing patterns that warrant further investigation into their potential role in recombination, introgression, and adaptation. Our findings reinforce the longstanding hypothesis that *Quercus* genomes exhibit a high degree of synteny, particularly between *Virentes* and its sister section *Quercus*. By expanding the genomic resources available for oaks, this assembly enhances our ability to investigate the evolutionary dynamics, adaptation, and hybridization potential within *Quercus*.

## Data Availability

The data used to generate these assemblies and annotations will be deposited on NCBI under BioProject PRJNA1310641. Custom scripts and methods used for the assembly, annotation and analyses are available at: https://github.com/laramiemckenna/quercus_virigniana_genome.

Supplementary data types and files will be available on Zenodo.

## Acknowledgements

We thank all donors to the Auburn University Tiger Giving Day project that funded the data generation of this project, as well as the Auburn University College of Agriculture and Department of Crop, Soil, and Environmental Science for co-funding the Praxis AI teaching platform expenses. A.H. was funded by National Science Foundation IOS-PGRP CAREER #2239530. L.A. was funded by National Science Foundation Graduate Research Fellowship Program #2239530.

## Author Contributions

**Concept and research design:** L.G., A.H.

**Sample collection, data collection, sequencing:** L.G., J.W., L.B.B., H.H., H.M., J.G., Z.M., A.H.

**Genome assembly and annotation:** L.A., Z.M., J.A., T.A., R.D., N.G., Z.P., S.P., G.R., C.A.R., A.H., C.B.

**Computational and statistical analysis:** L.A., Z.M., L.G., J.A.-M., T.A., R.D., N.G., Z.P., S.P., G.R., C.A.R., C.B., A.K., E.M., S.E.

**First draft writing and figure development:** L.A., Z.M., J.A.-M., T.A., R.D., N.G., Z.P., S.P. G.R., C.A.R., F.A.F., D.J.C.IV, J.S., J.L.

**Final draft writing and figure development (with contributions from all authors):** L.A., Z.M., S.B.C., L.G., A.H.

## References

Ai W, Liu Y, Mei M, Zhang X, Tan E, Liu H, Han X, Zhan H, Lu X. 2022. A chromosome-scale genome assembly of the Mongolian oak (Quercus mongolica). Mol Ecol Resour. 22(6):2396–2410. doi:10.1111/1755-0998.13616. https://onlinelibrary.wiley.com/doi/10.1111/1755-0998.13616.

Astashyn A, Tvedte ES, Sweeney D, Sapojnikov V, Bouk N, Joukov V, Mozes E, Strope PK, Sylla PM, Wagner L, et al. 2024. Rapid and sensitive detection of genome contamination at scale with FCS-GX. Genome Biol. 25(1):60. doi:10.1186/s13059-024-03198-7. https://genomebiology.biomedcentral.com/articles/10.1186/s13059-024-03198-7.

Benson G. 1999. Tandem repeats finder: a program to analyze DNA sequences. Nucleic Acids Res. 27(2):573–580. doi:10.1093/nar/27.2.573. https://pubmed.ncbi.nlm.nih.gov/9862982/.

Berdan EL, Aubier TG, Cozzolino S, Faria R, Feder JL, Giménez MD, Joron M, Searle JB, Mérot C. 2024. Structural variants and speciation: Multiple processes at play. Cold Spring Harb Perspect Biol. 16(3):a041446. doi:10.1101/cshperspect.a041446. http://dx.doi.org/10.1101/cshperspect.a041446.

Bradnam KR, Fass JN, Alexandrov A, Baranay P, Bechner M, Birol I, Boisvert S, Chapman JA, Chapuis G, Chikhi R, et al. 2013. Assemblathon 2: evaluating de novo methods of genome assembly in three vertebrate species. Gigascience. 2(1):10. doi:10.1186/2047-217X-2-10. https://academic.oup.com/gigascience/article/2/1/2047-217X-2-10/2656129.

Bushnell B. 2014 Mar 17. BBMap: A fast, accurate, splice-aware aligner. [accessed 2025 Feb 15]. https://www.osti.gov/servlets/purl/1241166.

Cannon CH, Kartesz J, Hoban S, Loza MI, Bruns EB, Hipp AL. 2024 Aug 7. Constructing sympatry networks to assess potential introgression pathways within the major oak sections in the contiguous US states. Plants People Planet. doi:10.1002/ppp3.10546. http://dx.doi.org/10.1002/ppp3.10546.

Cannon CH, Petit RJ. 2020. The oak syngameon: more than the sum of its parts. New Phytol. 226(4):978–983. doi:10.1111/nph.16091. http://dx.doi.org/10.1111/nph.16091.

Cavender-Bares J. 2019. Diversification, adaptation, and community assembly of the American oaks (Quercus), a model clade for integrating ecology and evolution. New Phytol. 221(2):669–692. doi:10.1111/nph.15450. https://nph.onlinelibrary.wiley.com/doi/10.1111/nph.15450.

Cavender-Bares J, González-Rodríguez A, Eaton DAR, Hipp AAL, Beulke A, Manos PS. 2015. Phylogeny and biogeography of the American live oaks (Quercus subsection Virentes): a genomic and population genetics approach. Mol Ecol. 24(14):3668–3687. doi:10.1111/mec.13269. http://dx.doi.org/10.1111/mec.13269.

Cheng H, Concepcion GT, Feng X, Zhang H, Li H. 2021. Haplotype-resolved de novo assembly using phased assembly graphs with hifiasm. Nat Methods. 18(2):170–175. doi:10.1038/s41592-020-01056-5. https://www.nature.com/articles/s41592-020-01056-5.

Chen N. 2004. Using RepeatMasker to identify repetitive elements in genomic sequences. Curr Protoc Bioinformatics. Chapter 4(1):Unit 4.10. doi:10.1002/0471250953.bi0410s05. https://pubmed.ncbi.nlm.nih.gov/18428725/.

Cuacos M, H Franklin FC, Heckmann S. 2015. Atypical centromeres in plants-what they can tell us. Front Plant Sci. 6:913. doi:10.3389/fpls.2015.00913. https://www.frontiersin.org/journals/plant-science/articles/10.3389/fpls.2015.00913/full.

De Coster W, D’Hert S, Schultz DT, Cruts M, Van Broeckhoven C. 2018. NanoPack: visualizing and processing long-read sequencing data. Bioinformatics. 34(15):2666–2669. doi:10.1093/bioinformatics/bty149. https://academic.oup.com/bioinformatics/article/34/15/2666/4934939.

Dupeyron M, Singh KS, Bass C, Hayward A. 2019. Evolution of Mutator transposable elements across eukaryotic diversity. Mob DNA. 10(1):12. doi:10.1186/s13100-019-0153-8. https://mobilednajournal.biomedcentral.com/articles/10.1186/s13100-019-0153-8.

Durand NC, Robinson JT, Shamim MS, Machol I, Mesirov JP, Lander ES, Aiden EL. 2016. Juicebox provides a visualization system for Hi-C contact maps with unlimited zoom. Cell Syst. 3(1):99–101. doi:10.1016/j.cels.2015.07.012. https://pmc.ncbi.nlm.nih.gov/articles/PMC5596920/.

Eaton DAR, Hipp AL, González-Rodríguez A, Cavender-Bares J. 2015. Historical introgression among the American live oaks and the comparative nature of tests for introgression. Evolution. 69(10):2587–2601. doi:10.1111/evo.12758. http://dx.doi.org/10.1111/evo.12758.

Elphinstone C, Elphinstone R, Todesco M, Rieseberg L. 2023. RepeatOBserver: tandem repeat visualization and centromere detection. Bioinformatics. https://www.biorxiv.org/content/10.1101/2023.12.30.573697v1.

Flynn JM, Hubley R, Goubert C, Rosen J, Clark AG, Feschotte C, Smit AF. 2020. RepeatModeler2 for automated genomic discovery of transposable element families. Proc Natl Acad Sci U S A. 117(17):9451–9457. doi:10.1073/pnas.1921046117. https://www.pnas.org/doi/10.1073/pnas.1921046117.

Gabriel L, Brůna T, Hoff KJ, Ebel M, Lomsadze A, Borodovsky M, Stanke M. 2023. BRAKER3: Fully automated genome annotation using RNA-seq and protein evidence with GeneMark-ETP, AUGUSTUS and TSEBRA. bioRxiv. doi:10.1101/2023.06.10.544449. https://pubmed.ncbi.nlm.nih.gov/37398387/.

Gao S, Jia Y, Guo H, Xu T, Wang B, Bush SJ, Wan S, Zhang Y, Yang X, Ye K. 2024. The centromere landscapes of four karyotypically diverse Papaver species provide insights into chromosome evolution and speciation. Cell Genom. 4(8):100626. doi:10.1016/j.xgen.2024.100626. https://www.cell.com/cell-genomics/fulltext/S2666-979X(24)00230-1.

Garg V, Bohra A, Mascher M, Spannagl M, Xu X, Bevan MW, Bennetzen JL, Varshney RK. 2024. Unlocking plant genetics with telomere-to-telomere genome assemblies. Nat Genet. 56(9):1788–1799. doi:10.1038/s41588-024-01830-7. https://www.nature.com/articles/s41588-024-01830-7.

Gel B, Serra E. 2017. karyoploteR: an R/Bioconductor package to plot customizable genomes displaying arbitrary data. Bioinformatics. 33(19):3088–3090. doi:10.1093/bioinformatics/btx346. https://academic.oup.com/bioinformatics/article/33/19/3088/3857734.

Goel M, Schneeberger K. 2022. Plotsr: Visualizing structural similarities and rearrangements between multiple genomes. Bioinformatics. 38(10):2922–2926. doi:10.1093/bioinformatics/btac196. https://academic.oup.com/bioinformatics/article/38/10/2922/6569079.

Goel M, Sun H, Jiao W-B, Schneeberger K. 2019. SyRI: finding genomic rearrangements and local sequence differences from whole-genome assemblies. Genome Biol. 20(1):277. doi:10.1186/s13059-019-1911-0. https://genomebiology.biomedcentral.com/articles/10.1186/s13059-019-1911-0.

Greiner S, Lehwark P, Bock R. 2019. OrganellarGenomeDRAW (OGDRAW) version 1.3.1: expanded toolkit for the graphical visualization of organellar genomes. Nucleic Acids Res. 47(W1):W59–W64. doi:10.1093/nar/gkz238. https://academic.oup.com/nar/article/47/W1/W59/5428289.

Han B, Wang L, Xian Y, Xie X-M, Li W-Q, Zhao Y, Zhang R-G, Qin X, Li D-Z, Jia K-H. 2022. A chromosome-level genome assembly of the Chinese cork oak (Quercus variabilis). Front Plant Sci. 13:1001583. doi:10.3389/fpls.2022.1001583. https://www.frontiersin.org/journals/plant-science/articles/10.3389/fpls.2022.1001583/full.

Hart AJ, Ginzburg S, Xu MS, Fisher CR, Rahmatpour N, Mitton JB, Paul R, Wegrzyn JL. 2020. EnTAP: Bringing faster and smarter functional annotation to non-model eukaryotic transcriptomes. Mol Ecol Resour. 20(2):591–604. doi:10.1111/1755-0998.13106. https://pubmed.ncbi.nlm.nih.gov/31628884/.

Hipp AL, Manos PS, Hahn M, Avishai M, Bodénès C, Cavender-Bares J, Crowl AA, Deng M, Denk T, Fitz-Gibbon S, et al. 2020. Genomic landscape of the global oak phylogeny. New Phytol. 226(4):1198–1212. doi:10.1111/nph.16162. https://nph.onlinelibrary.wiley.com/doi/10.1111/nph.16162.

Hipp AL, Whittemore AT, Garner M, Hahn M, Fitzek E, Guichoux E, Cavender-Bares J, Gugger PF, Manos PS, Pearse IS, et al. 2019. Genomic identity of White Oak species in an eastern North American syngameon. Ann Mo Bot Gard. 104(3):455–477. doi:10.3417/2019434. https://bioone.org/journals/annals-of-the-missouri-botanical-garden/volume-104/issue-3/2019434/----Custom-HTMLGenomic/10.3417/2019434.short.

Huang G, Bao Z, Feng L, Zhai J, Wendel JF, Cao X, Zhu Y. 2024. A telomere-to-telomere cotton genome assembly reveals centromere evolution and a Mutator transposon-linked module regulating embryo development. Nat Genet. 56(9):1953–1963. doi:10.1038/s41588-024-01877-6. https://www.nature.com/articles/s41588-024-01877-6.

Huang N, Li H. 2023. compleasm: a faster and more accurate reimplementation of BUSCO. Bioinformatics. 39(10):btad595. doi:10.1093/bioinformatics/btad595. https://academic.oup.com/bioinformatics/article/39/10/btad595/7284108.

Hu H-L, Zhang J-Y, Li Y-P, Xie L, Chen D-B, Li Q, Liu Y-Q, Hui S-R, Qin L. 2019. The complete chloroplast genome of the daimyo oak, Quercus dentata Thunb. Conserv Genet Resour. 11(4):409–411. doi:10.1007/s12686-018-1034-z. https://link.springer.com/article/10.1007/s12686-018-1034-z.

International Oak Society. Date unknown. Hybrid Highlight: Quercus ×comptoniae Sarg. [Internet]; [cited 2025 Aug 19]. Available from: https://www.internationaloaksociety.org/content/hybrid-highlight-quercus-%C3%97comp toniae-sarg.

Kapoor B, Jenkins J, Schmutz J, Zhebentyayeva T, Kuelheim C, Coggeshall M, Heim C, Lasky JR, Leites L, Islam-Faridi N, et al. 2023. A haplotype-resolved chromosome-scale genome for Quercus rubra L. provides insights into the genetics of adaptive traits for red oak species. G3 (Bethesda). 13(11):jkad209. doi:10.1093/g3journal/jkad209. https://academic.oup.com/g3journal/article/13/11/jkad209/7274082.

Kremer A, Hipp AL. 2020. Oaks: an evolutionary success story. New Phytol. 226(4):987–1011. doi:10.1111/nph.16274. http://dx.doi.org/10.1111/nph.16274.

Larson DA, Staton ME, Kapoor B, Islam-Faridi N, Zhebentyayeva T, Fan S, Stork J, Thomas A, Ahmed AS, Stanton EC, et al. 2025 Feb 11. A haplotype-resolved reference genome of Quercus alba sheds light on the evolutionary history of oaks. New Phytol. doi:10.1111/nph.20463. http://dx.doi.org/10.1111/nph.20463.

Li H, Durbin R. 2009. Fast and accurate short read alignment with Burrows-Wheeler transform. Bioinformatics. 25(14):1754–1760. doi:10.1093/bioinformatics/btp324. https://academic.oup.com/bioinformatics/article/25/14/1754/225615.

Liu Y, Yi C, Fan C, Liu Q, Liu S, Shen L, Zhang K, Huang Y, Liu C, Wang Y, et al. 2023. Pan-centromere reveals widespread centromere repositioning of soybean genomes. Proc Natl Acad Sci U S A. 120(42):e2310177120. doi:10.1073/pnas.2310177120. https://www.pnas.org/doi/epub/10.1073/pnas.2310177120.

Lovell JT, Sreedasyam A, Schranz ME, Wilson M, Carlson JW, Harkess A, Emms D, Goodstein DM, Schmutz J. 2022. GENESPACE tracks regions of interest and gene copy number variation across multiple genomes. Elife. 11:e78526. doi:10.7554/eLife.78526. https://elifesciences.org/articles/78526.

Luo C-S, Li T-T, Jiang X-L, Song Y, Fan T-T, Shen X-B, Yi R, Ao X-P, Xu G-B, Deng M. 2024. High-quality haplotype-resolved genome assembly for ring-cup oak (Quercus glauca) provides insight into oaks demographic dynamics. Mol Ecol Resour. 24(3):e13914. doi:10.1111/1755-0998.13914. https://onlinelibrary.wiley.com/doi/10.1111/1755-0998.13914.

Macas J, Ávila Robledillo L, Kreplak J, Novák P, Koblížková A, Vrbová I, Burstin J, Neumann P. 2023. Assembly of the 81.6 Mb centromere of pea chromosome 6 elucidates the structure and evolution of metapolycentric chromosomes. PLoS Genet. 19(2):e1010633. doi:10.1371/journal.pgen.1010633. https://journals.plos.org/plosgenetics/article?id=10.1371/journal.pgen.1010633.

Ma L, Li G. 2018. FAR1-RELATED SEQUENCE (FRS) and FRS-RELATED FACTOR (FRF) family proteins in Arabidopsis growth and development. Front Plant Sci. 9:692. doi:10.3389/fpls.2018.00692. https://www.frontiersin.org/journals/plant-science/articles/10.3389/fpls.2018.00692/full#B29.

Manni M, Berkeley MR, Seppey M, Zdobnov EM. 2021. BUSCO: Assessing genomic data quality and beyond. Curr Protoc. 1(12):e323. doi:10.1002/cpz1.323. https://currentprotocols.onlinelibrary.wiley.com/doi/full/10.1002/cpz1.323.

McVay JD, Hipp AL, Manos PS. 2017. A genetic legacy of introgression confounds phylogeny and biogeography in oaks. Proc Biol Sci. 284(1854). doi:10.1098/rspb.2017.0300. https://royalsocietypublishing.org/doi/10.1098/rspb.2017.0300.

Mead A, Fitz-Gibbon ST, Escalona M, Beraut E, Sacco S, Marimuthu MPA, Nguyen O, Sork VL. 2024. The genome assembly of Island Oak (Quercus tomentella), a relictual island tree species. J Hered. 115(2):221–229. doi:10.1093/jhered/esae002. https://academic.oup.com/jhered/article/115/2/221/7596578.

Naish M. 2024. Bridging the gap: unravelling plant centromeres in the telomere-to-telomere era. New Phytol. 244(6):2143–2149. doi:10.1111/nph.20149. http://dx.doi.org/10.1111/nph.20149.

Naish M, Henderson IR. 2024. The structure, function, and evolution of plant centromeres. Genome Res. 34(2):161–178. doi:10.1101/gr.278409.123. http://dx.doi.org/10.1101/gr.278409.123.

Nesom GL. 2018. Quercus × harbsonii: a naturally occurring hybrid in Texas and Oklahoma [Internet]; [cited 2025 Aug 19]. Available from: https://bugwoodcloud.org/resource/files/18742.pdf.

Open2C, Abdennur N, Fudenberg G, Flyamer IM, Galitsyna AA, Goloborodko A, Imakaev M, Venev SV. 2024. Pairtools: From sequencing data to chromosome contacts. PLoS Comput Biol. 20(5):e1012164. doi:10.1371/journal.pcbi.1012164. https://journals.plos.org/ploscompbiol/article?id=10.1371/journal.pcbi.1012164.

Ou S, Su W, Liao Y, Chougule K, Agda JRA, Hellinga AJ, Lugo CSB, Elliott TA, Ware D, Peterson T, et al. 2019. Benchmarking transposable element annotation methods for creation of a streamlined, comprehensive pipeline. Genome Biol. 20(1):275. doi:10.1186/s13059-019-1905-y. https://genomebiology.biomedcentral.com/articles/10.1186/s13059-019-1905-y.

Plomion C, Aury J-M, Amselem J, Leroy T, Murat F, Duplessis S, Faye S, Francillonne N, Labadie K, Le Provost G, et al. 2018. Oak genome reveals facets of long lifespan. Nat Plants. 4(7):440–452. doi:10.1038/s41477-018-0172-3. https://www.nature.com/articles/s41477-018-0172-3.

Ranallo-Benavidez TR, Rhyker Ranallo-Benavidez T, Jaron KS, Schatz MC. 2020. GenomeScope 2.0 and Smudgeplot for reference-free profiling of polyploid genomes. Nature Communications. 11(1). doi:10.1038/s41467-020-14998-3. http://dx.doi.org/10.1038/s41467-020-14998-3.

Rey M-D, Labella-Ortega M, Guerrero-Sánchez VM, Carleial R, Castillejo MÁ, Ruggieri V, Jorrín-Novo JV. 2023. A first draft genome of holm oak (Quercus ilex subsp. ballota), the most representative species of the Mediterranean forest and the Spanish agrosylvopastoral ecosystem “dehesa.” Front Mol Biosci. 10:1242943. doi:10.3389/fmolb.2023.1242943. https://www.frontiersin.org/journals/molecular-biosciences/articles/10.3389/fmolb.2023.1242943/full.

Rhie A, Walenz BP, Koren S, Phillippy AM. 2020. Merqury: reference-free quality, completeness, and phasing assessment for genome assemblies. Genome Biol. 21(1):245. doi:10.1186/s13059-020-02134-9. https://genomebiology.biomedcentral.com/articles/10.1186/s13059-020-02134-9.

Rieseberg LH. 2001. Chromosomal rearrangements and speciation. Trends Ecol Evol. 16(7):351–358. doi:10.1016/s0169-5347(01)02187-5. http://dx.doi.org/10.1016/s0169-5347(01)02187-5.

Sandbox Bio. dot: an interactive dot plot viewer for genome-genome alignments [Internet]. [cited 2025 Aug 26]. Available from: https://dot.sandbox.bio/

Sork VL, Cokus SJ, Fitz-Gibbon ST, Zimin AV, Puiu D, Garcia JA, Gugger PF, Henriquez CL, Zhen Y, Lohmueller KE, et al. 2022. High-quality genome and methylomes illustrate features underlying evolutionary success of oaks. Nat Commun. 13(1):2047. doi:10.1038/s41467-022-29584-y. https://www.nature.com/articles/s41467-022-29584-y.

Tillich M, Lehwark P, Pellizzer T, Ulbricht-Jones ES, Fischer A, Bock R, Greiner S. 2017. GeSeq - versatile and accurate annotation of organelle genomes. Nucleic Acids Res. 45(W1):W6–W11. doi:10.1093/nar/gkx391. https://academic.oup.com/nar/article/45/W1/W6/3806659.

Vaser R, Sović I, Nagarajan N, Šikić M. 2017. Fast and accurate de novo genome assembly from long uncorrected reads. Genome Res. 27(5):737–746. doi:10.1101/gr.214270.116. https://pmc.ncbi.nlm.nih.gov/articles/PMC5411768/.

Vollger MR, Kerpedjiev P, Phillippy AM, Eichler EE. 2022. StainedGlass: interactive visualization of massive tandem repeat structures with identity heatmaps. Bioinformatics. 38(7):2049–2051. doi:10.1093/bioinformatics/btac018. https://academic.oup.com/bioinformatics/article/38/7/2049/6502275.

Wang G, Li H, Cheng Z, Jin W. 2013. A novel translocation event leads to a recombinant stable chromosome with interrupted centromeric domains in rice. Chromosoma. 122(4):295–303. doi:10.1007/s00412-013-0413-1. http://dx.doi.org/10.1007/s00412-013-0413-1.

Wang W-B, He X-F, Yan X-M, Ma B, Lu C-F, Wu J, Zheng Y, Wang W-H, Xue W-B, Tian X-C, et al. 2023. Chromosome-scale genome assembly and insights into the metabolome and gene regulation of leaf color transition in an important oak species, Quercus dentata. New Phytol. 238(5):2016–2032. doi:10.1111/nph.18814. https://nph.onlinelibrary.wiley.com/doi/full/10.1111/nph.18814.

Wellcome Sanger Institute. 2022. Quercus robur genome assembly dhQueRobu3.1. [dataset]. [Internet]. Bethesda (MD): NCBI GenBank; [cited 2025 Aug 18]. Available from: https://www.ncbi.nlm.nih.gov/datasets/genome/GCF_932294415.1/.

Whittemore AT, Miller RE. 2023. Dynamic properties of the pinyon pine syngameon. New Phytol. 237(6):1943–1945. doi:10.1111/nph.18707. https://nph.onlinelibrary.wiley.com/doi/10.1111/nph.18707.

Wlodzimierz P, Hong M, Henderson IR. 2023. TRASH: Tandem Repeat Annotation and Structural Hierarchy. Bioinformatics. 39(5):btad308. doi:10.1093/bioinformatics/btad308. https://academic.oup.com/bioinformatics/article/39/5/btad308/7159186?login=false.

Xian W, Carbonell-Bejerano P, Rabanal FA, Bezrukov I, Reymond P, Weigel D. 2025. Minimizing detection bias of somatic mutations in a highly heterozygous oak genome. Genomics. https://www.biorxiv.org/content/10.1101/2025.02.13.638107v1.

Yang Y, Zhou T, Duan D, Yang J, Feng L, Zhao G. 2016. Comparative analysis of the complete chloroplast genomes of five Quercus species. Front Plant Sci. 7:959. doi:10.3389/fpls.2016.00959. https://www.frontiersin.org/journals/plant-science/articles/10.3389/fpls.2016.00959/full.

Zhang R-S, Yang J, Hu H-L, Xia R-X, Li Y-P, Su J-F, Li Q, Liu Y-Q, Qin L. 2020. A high level of chloroplast genome sequence variability in the Sawtooth Oak Quercus acutissima. Int J Biol Macromol. 152:340–348. doi:10.1016/j.ijbiomac.2020.02.201. https://www.sciencedirect.com/science/article/pii/S0141813019388841?casa_token=3Zc4AdN_OJYAAAAA:MaXx7ub5iCKvdaxwcm33nPWQd4GWiZTYNNvLr74GS8L-zx35C8fOi0ox45mDyVk6WUZO3xYEeo8.

Zhou C, Brown M, Blaxter M, McCarthy SA, Durbin R. 2024. Oatk: a de novo assembly tool for complex plant organelle genomes. Bioinformatics. https://www.biorxiv.org/content/10.1101/2024.10.23.619857v1.

Zhou C, McCarthy SA, Durbin R. 2023. YaHS: yet another Hi-C scaffolding tool. Bioinformatics. 39(1):btac808. doi:10.1093/bioinformatics/btac808. https://academic.oup.com/bioinformatics/article/39/1/btac808/6917071.

Zhou X, Liu N, Jiang X, Qin Z, Farooq TH, Cao F, Li H. 2022. A chromosome-scale genome assembly of Quercus gilva: Insights into the evolution of Quercus section Cyclobalanopsis (Fagaceae). Front Plant Sci. 13:1012277. doi:10.3389/fpls.2022.1012277. https://www.frontiersin.org/journals/plant-science/articles/10.3389/fpls.2022.1012277/full.

