## Supplementary Material for "A haplotype-resolved, chromosome-scale genome assembly for the southern live oak, *Quercus virginiana*"

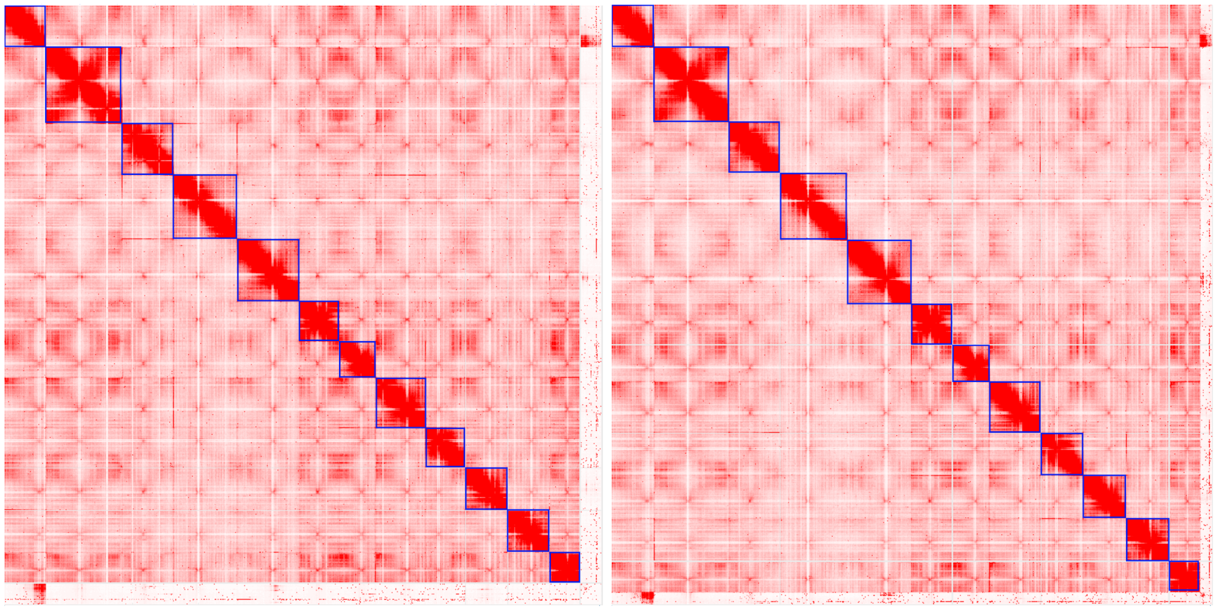

**Fig. S1. Contact maps after manual curation.** Haplotype 1 is represented on the left, and Haplotype 2 is represented on the right. Contacts suggest that parts of Chromosome 1 in both haplotypes may interact with elements in the debris bin. However, these elements were revealed to be highly fragmented and unresolved repeats. They remain in the assembly as unincorporated scaffolds.

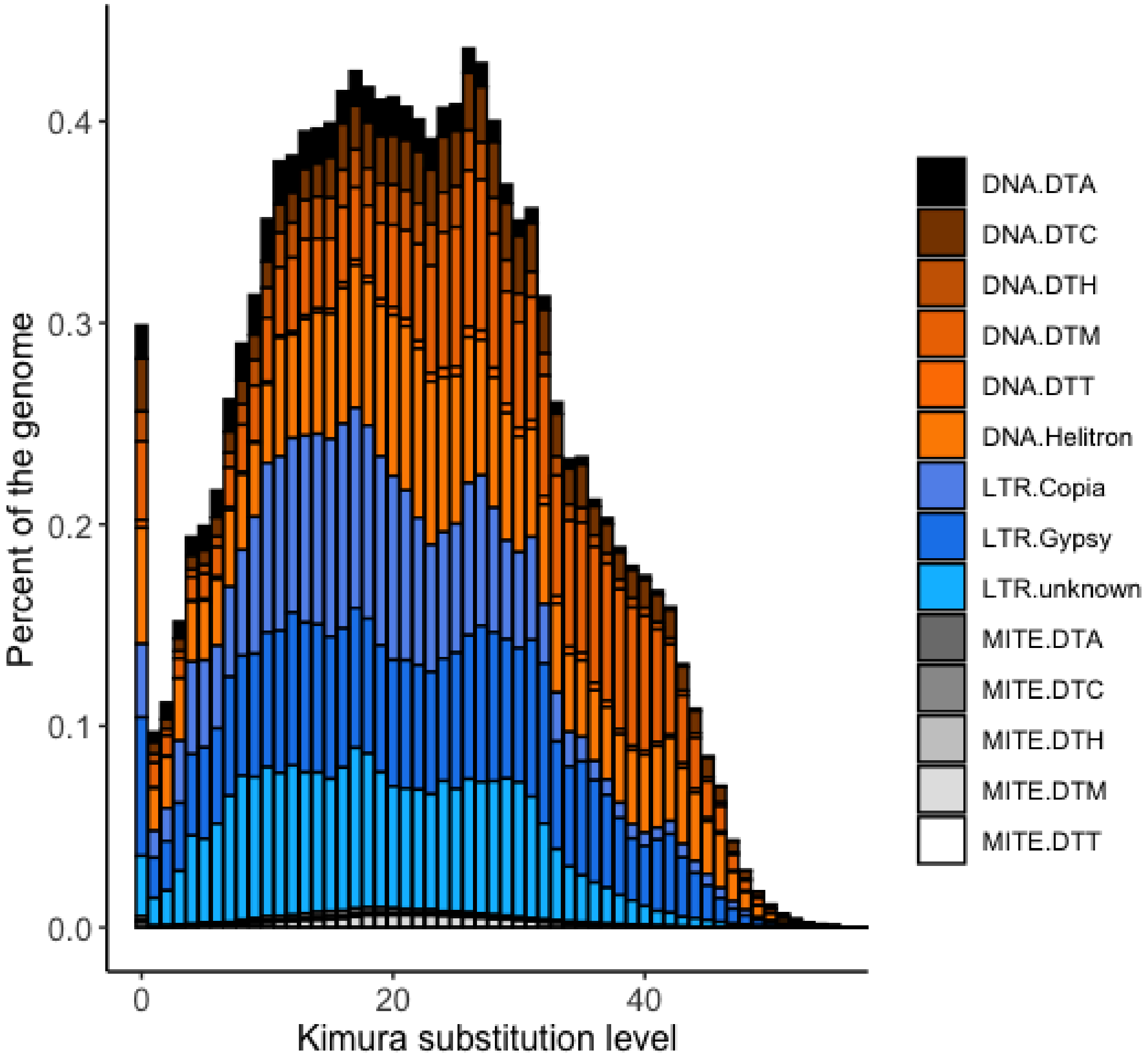


**Fig. S2. TE integration into *Quercus virginiana*.** Relative time was determined using the Kimura substitution level, with lower values closer to 0 representing more recent events and higher values approaching 40 representing older events, showing a burst in the introduction of TEs into the genome at roughly 1, 18, and 26 on the x-axis.

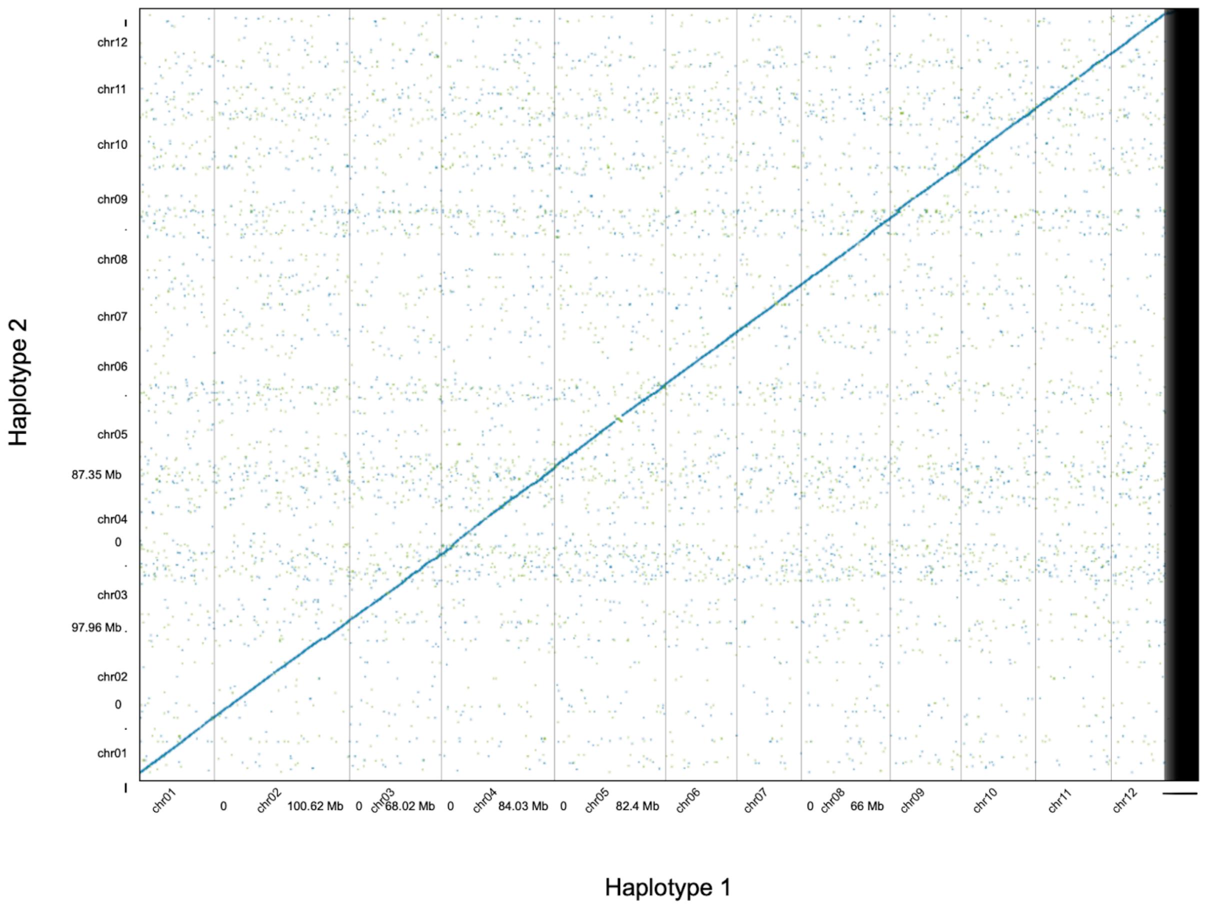


**Fig. S3. Dotplot comparing the haplotypes of *Q. virginiana*.** Comparisons of the haplotypes with an all-by-all alignment with *nucmer* reveal minimal structural variation between the the haplotypes except at putatively centromeric locations on Chr03, Chr05, Chr07 and Chr11.

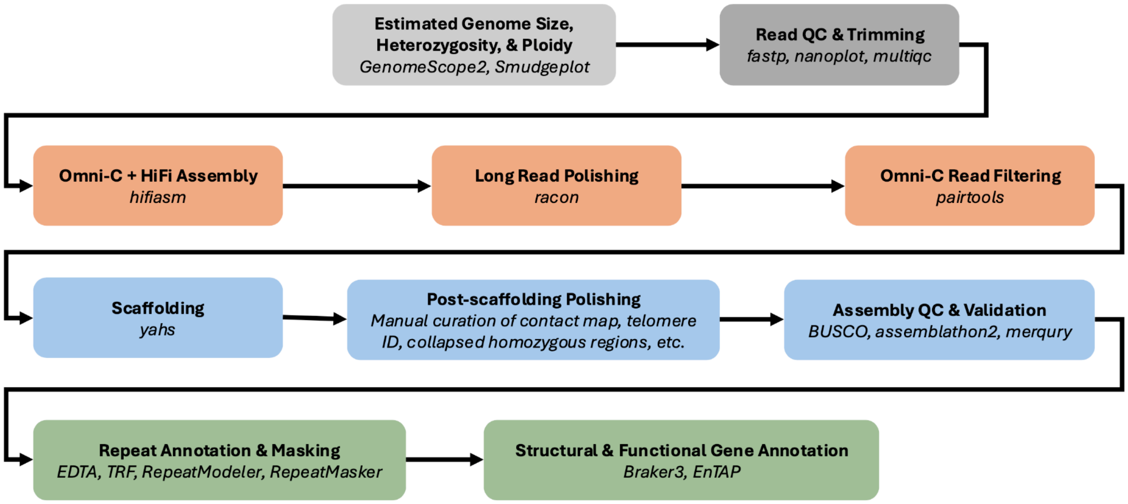


**Fig. S4. Genome assembly workflow.** Prior to beginning assembly, the *Q. virginiana* genome was first assessed for its genome size, heterozygosity and ploidy with Illumina reads. Later, the HiFi and Omni-C reads were checked for their quality. After an initial contig-level assembly with *hifiasm*, haplotype-aware long-read polishing was performed with *racon* and Omni-C reads were filtered for duplicates, sorted, and unmapped reads were removed with *pairtools* prior to scaffolding with *yahs*. Post-scaffolding polishing steps included assessing telomere location, checking for collapsed homozygous regions, manually curating the contact map, and more. Final quality checks and chromosome naming as well as orientation preceded structural gene annotation with *Braker3*, transposable element annotation and tandem repeat annotation with *EDTA, RepeatModeler* and *TRF*, and functional annotation with *EnTAP*.

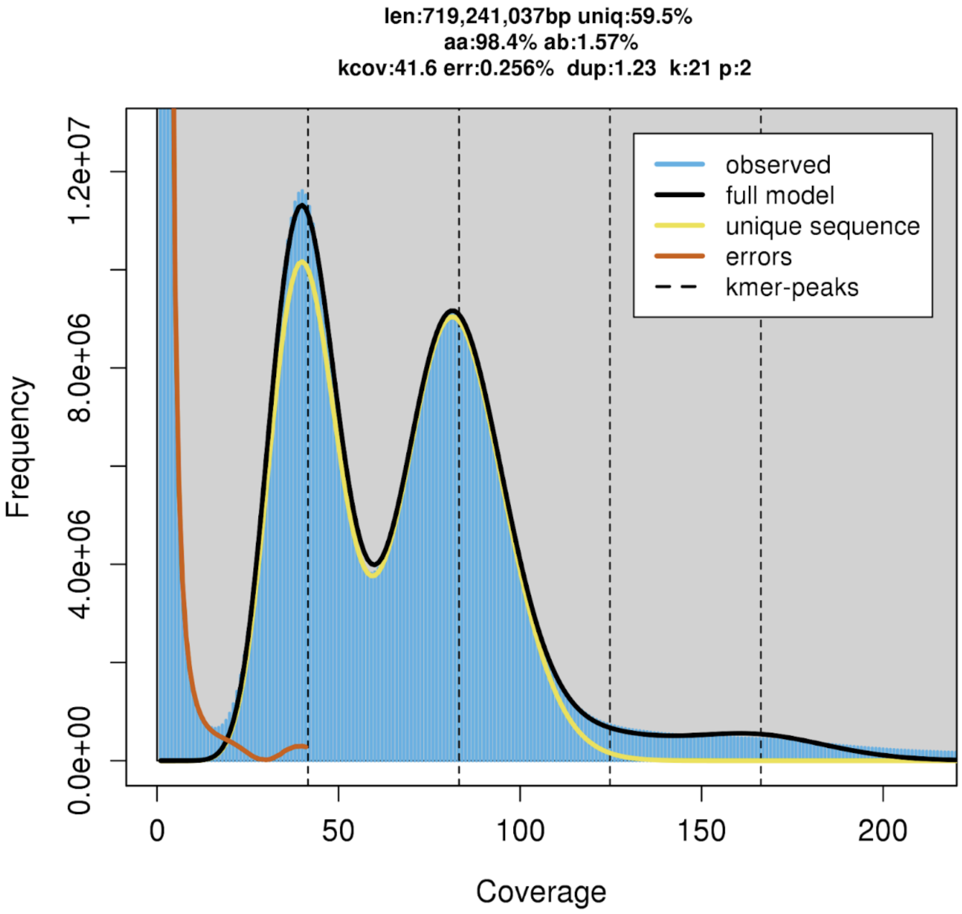


**Fig. S5. K-mer frequency plot for *Q. virginiana.*** Generated by GenomeScope2 at a k-mer length of k =21. The peak on the right side is the diploid/homozygous peak at a depth of 83 and the peak on the left side is the haploid/heterozygous peak at a depth of 41.5. The computed length for this genome after the removal of sequencing errors was 719Mb. The nucleotides had 1.57% heterozygosity. The error rate was 0.256% and the PCR duplication rate was 1.23%.

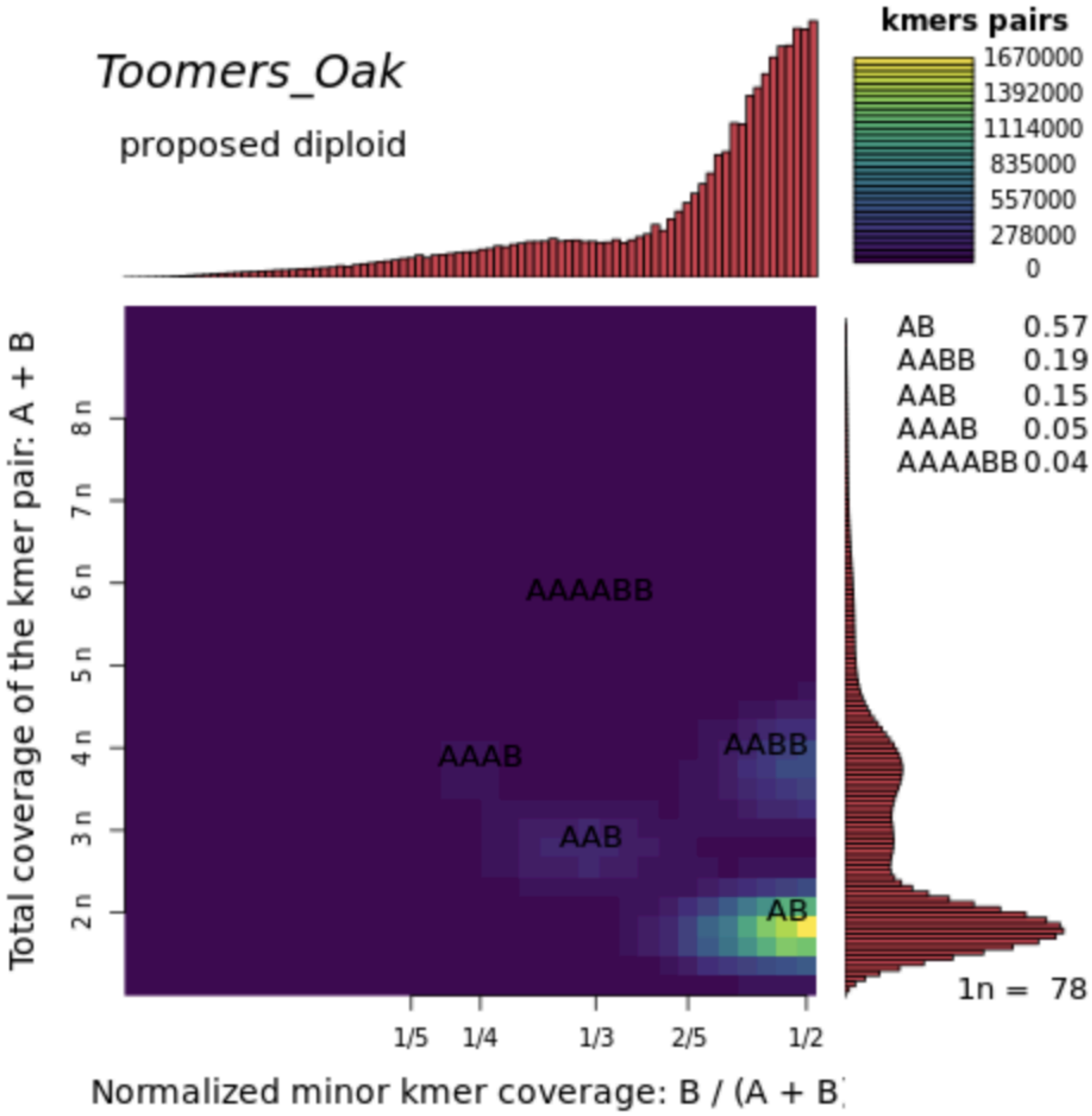


**Fig. S6. Smudgeplot describing ploidy of the *Q. virginiana* genome.** SmudgePlot showing over 57% of the k-mers are AB proposing *Quercus virginiana* is diploid. Peaks are split up into AB, AABB, AAB, AAAB, and AAAABB.

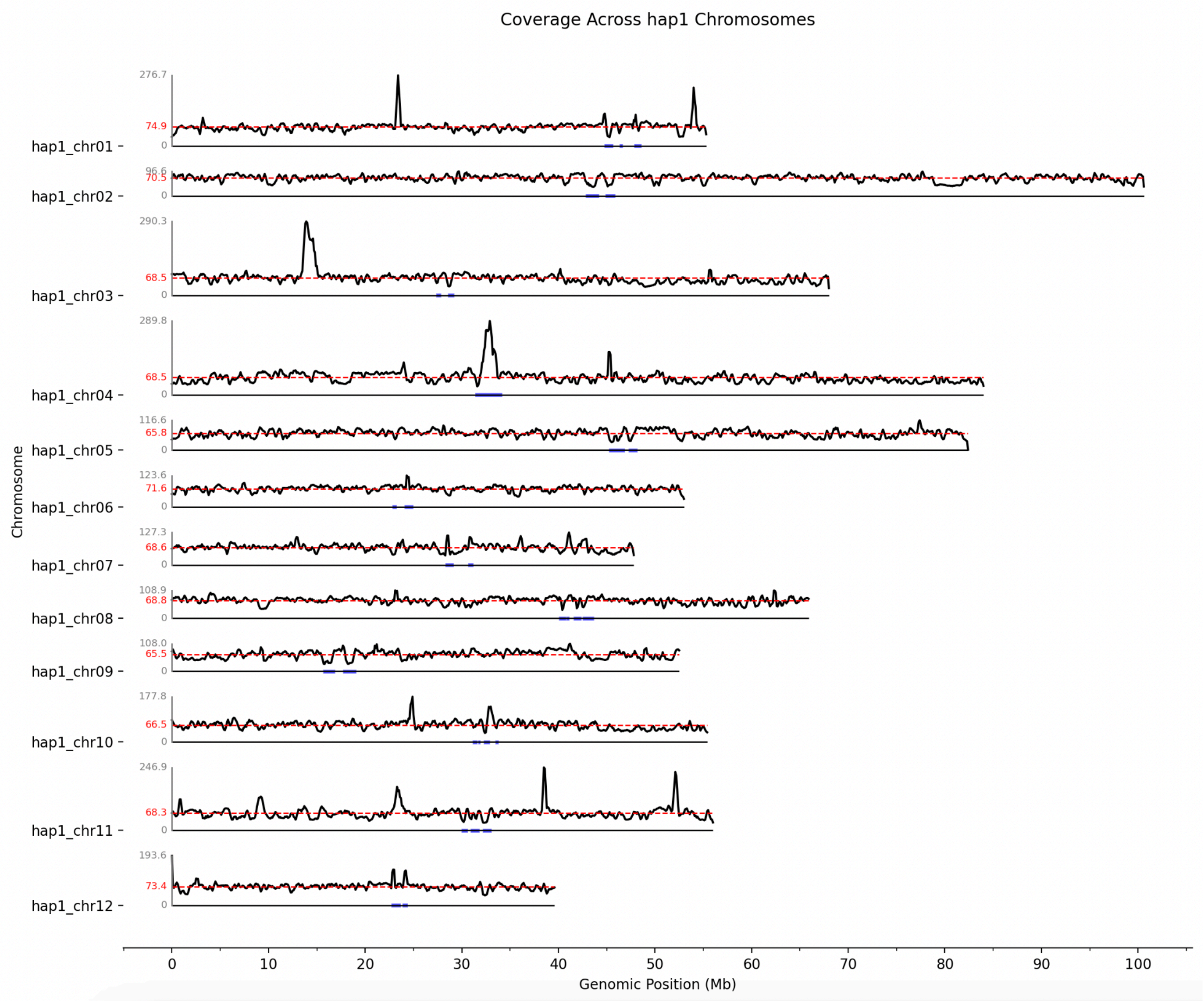
Coverage Across hap1 Chromosomes
Chromosome
hap1_chr01
hap1_chr02
hap1_chr03
hap1_chr04
hap1_chr05
hap1_chr06
hap1_chr07
hap1_chr08
hap1_chr09
hap1_chr10
hap1_chr11
hap1_chr12
Genomic Position (Mb)

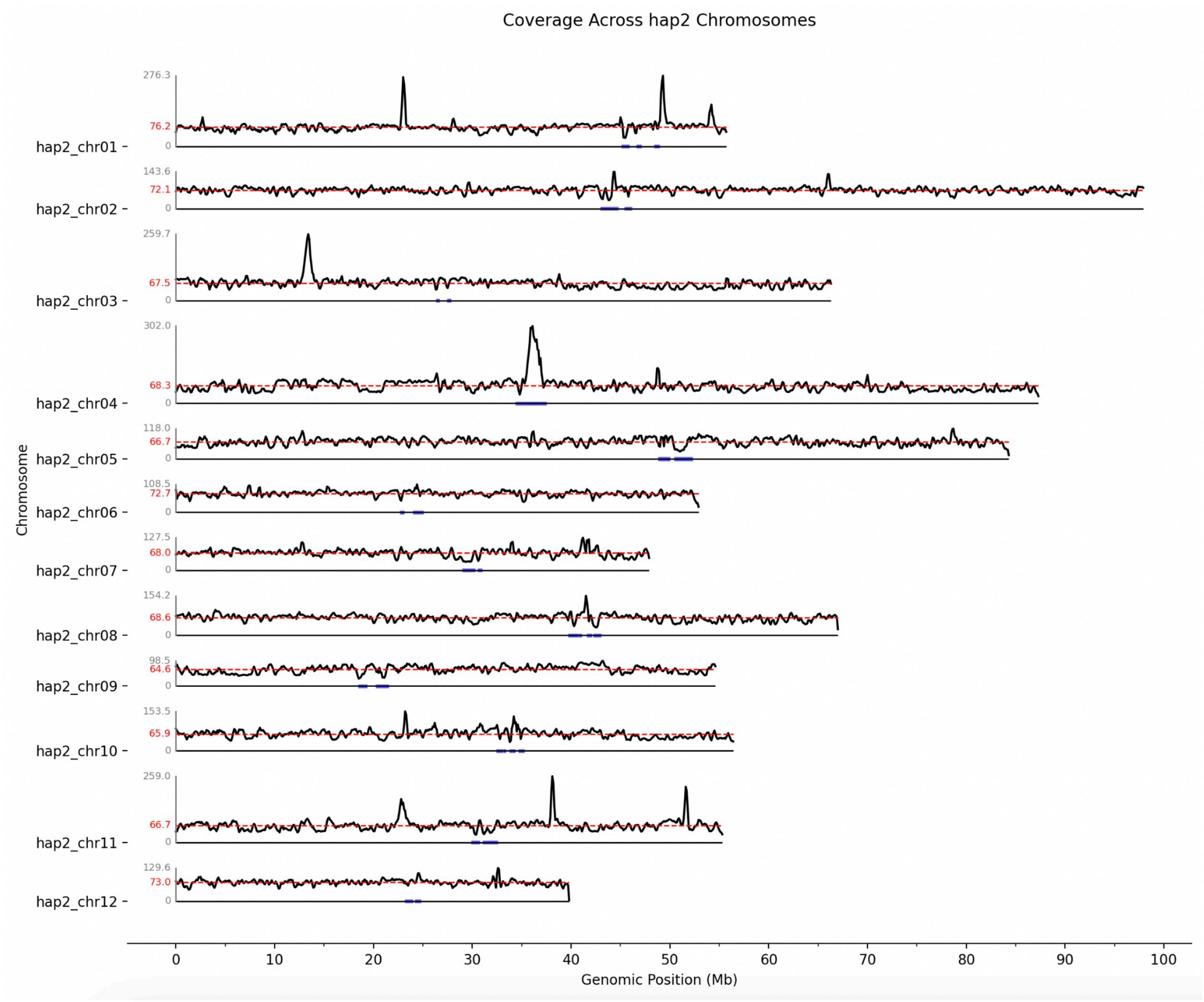


**Fig. S7. Density of HiFi reads along each haplotype at 300kb windows in 100kb increments.** Genomic position is represented in megabasepairs on the x-axis. For each chromosome, the average density and maximum density values are represented on the y-axis. Coverage is relatively even across the genome with some regions indicating higher or lower levels of coverage relative to the average. These regions correspond to repeat arrays. Excessive coverage may indicate a collapsed region or sequencing bias.

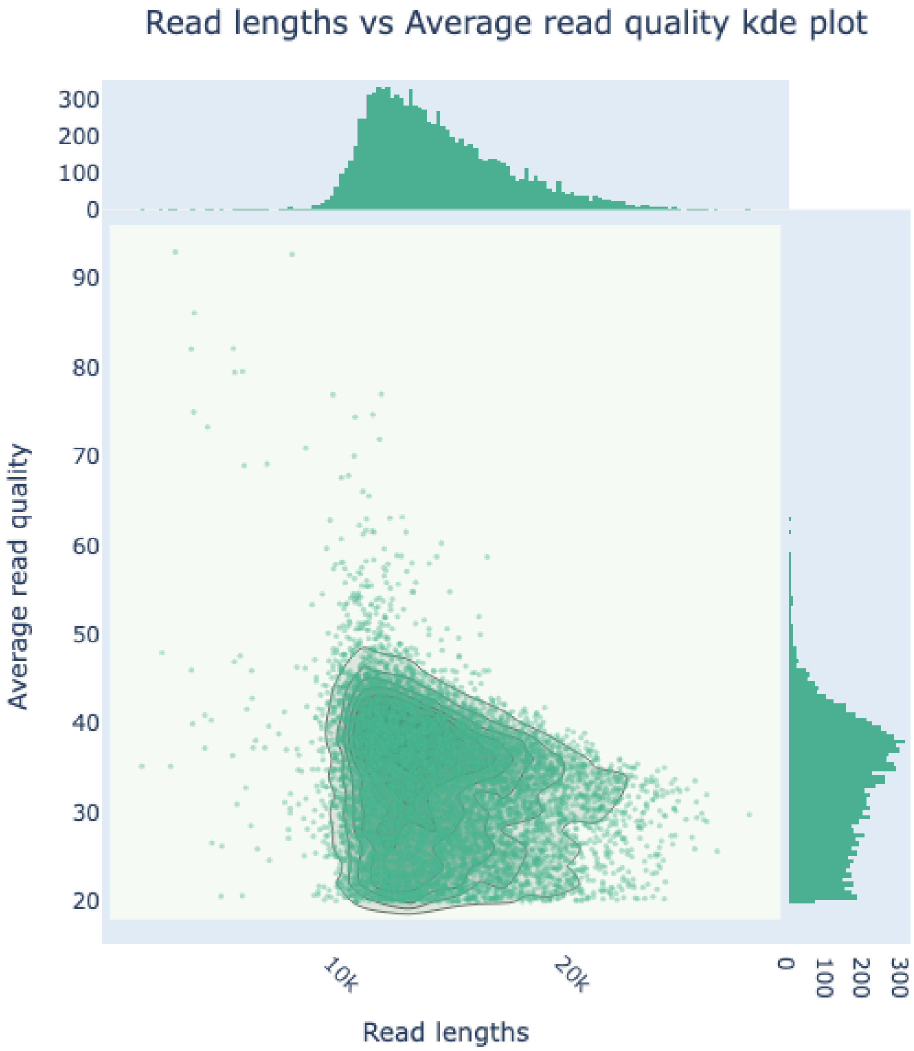


**Fig. S8. HiFi read length and quality.** Over 4.56 million HiFi reads were generated, totalling approximately 66.46 billion bases with a mean read length of 14,570 bp. 100% of the reads generated were >Q15 and the majority of reads exceeded 10,000 bp.

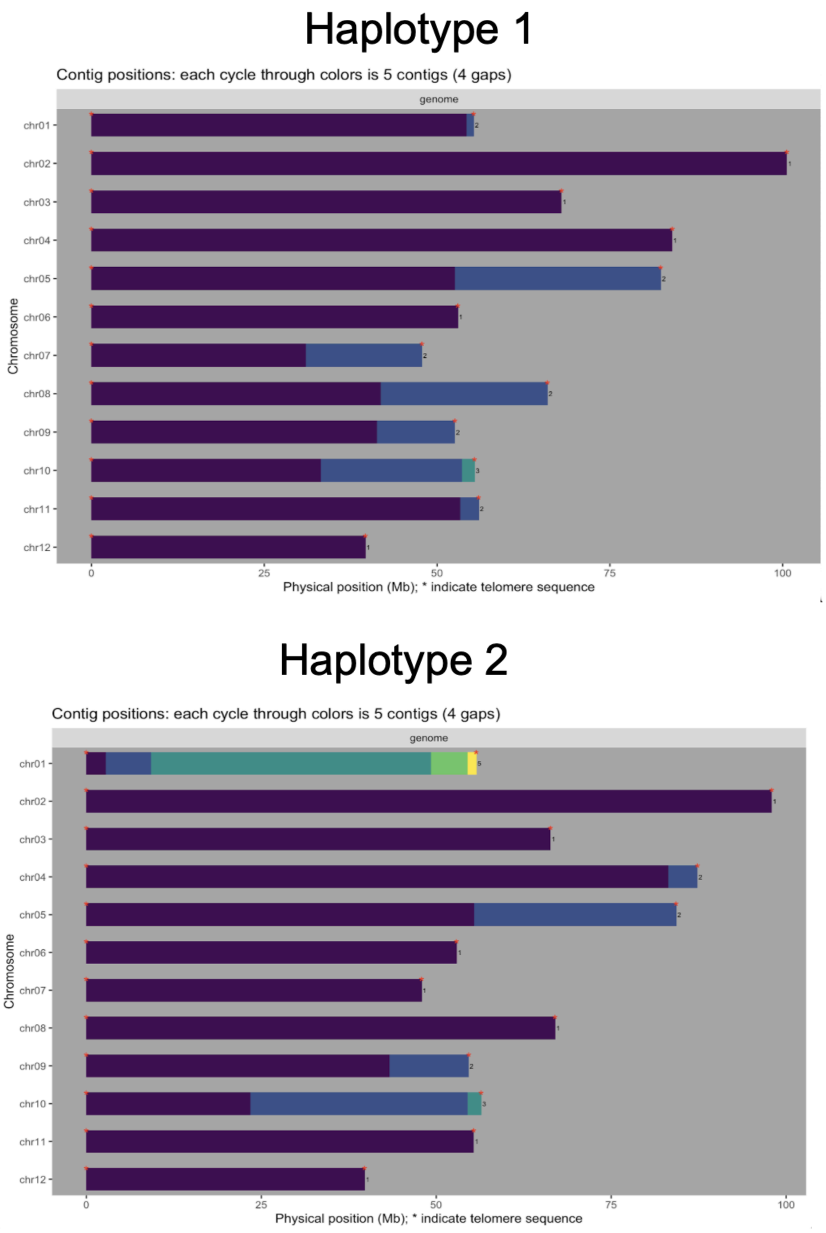


**Fig. S9. Assessing contiguity and telomere presence/absence.** GENESPACE was used to assess the contiguity of the *Q. virginiana* haplotypes. This genome is characterized by high contiguity and minimal gaps, with only eight gaps reported in Haplotype 1 and nine in Haplotype 2. Most chromosomes required minimal scaffolding, as the entire chromosome was represented in an entire contig. Telomeric sequence was identified at the ends of all chromosomes across both haplotypes.

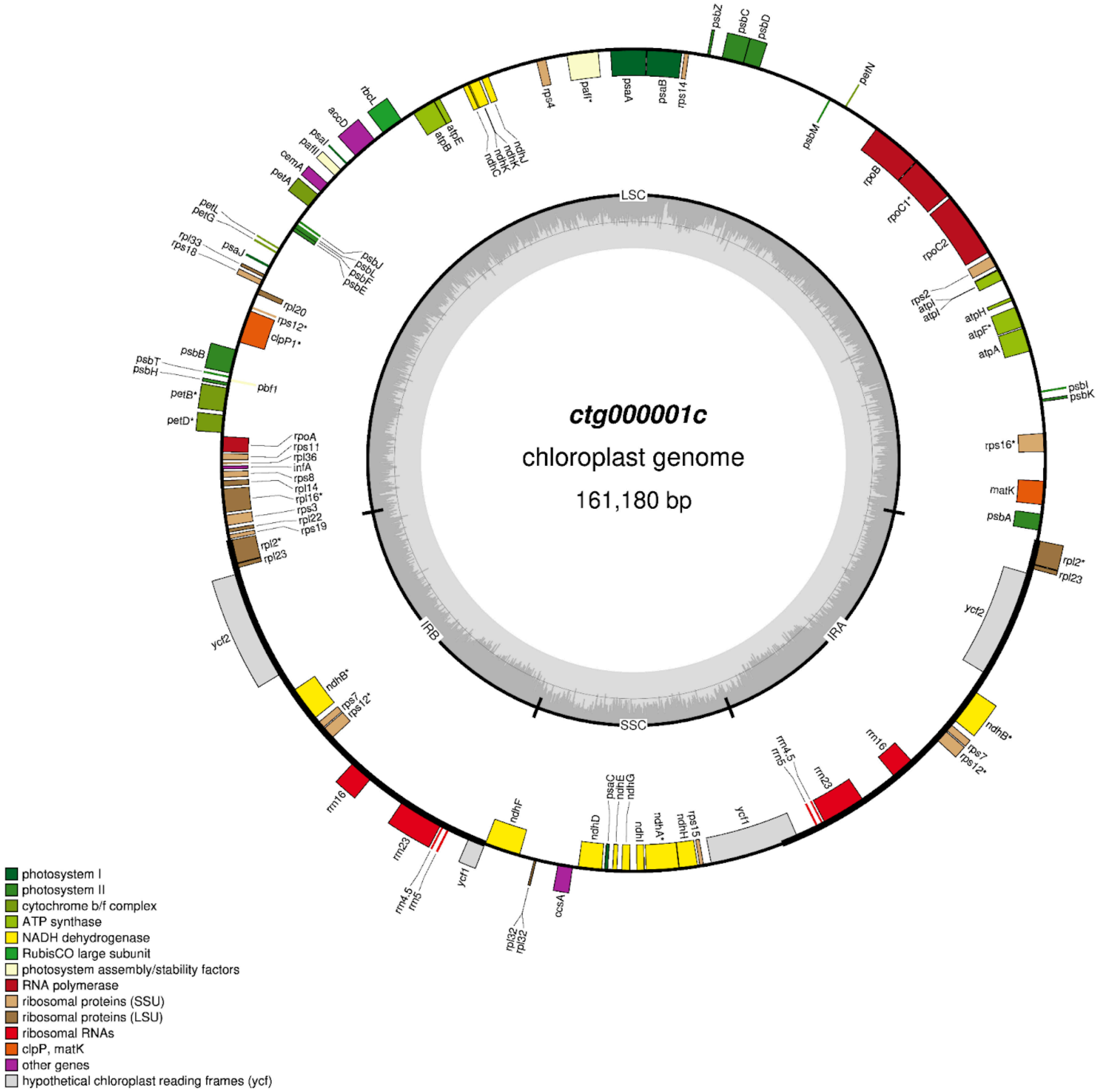


**Fig. S10, The *Q. virginiana* chloroplast assembly.** *OatK* was used to generate the 161,180 bp *Q. virginiana* chloroplast genome from HiFi reads.

### Haplotype 1 Centromeric Locations

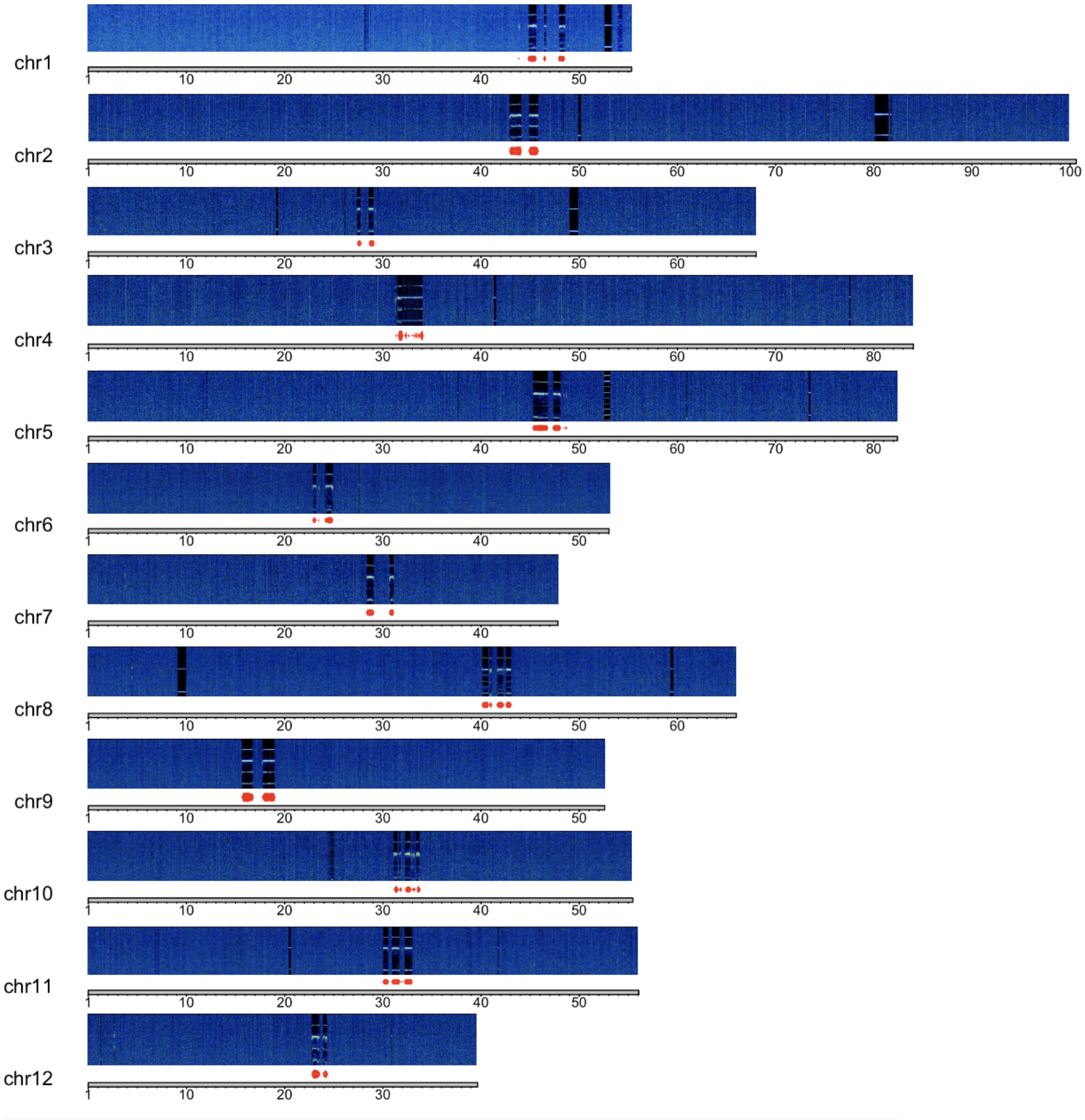

**Haplotype 2 Centromeric Locations**

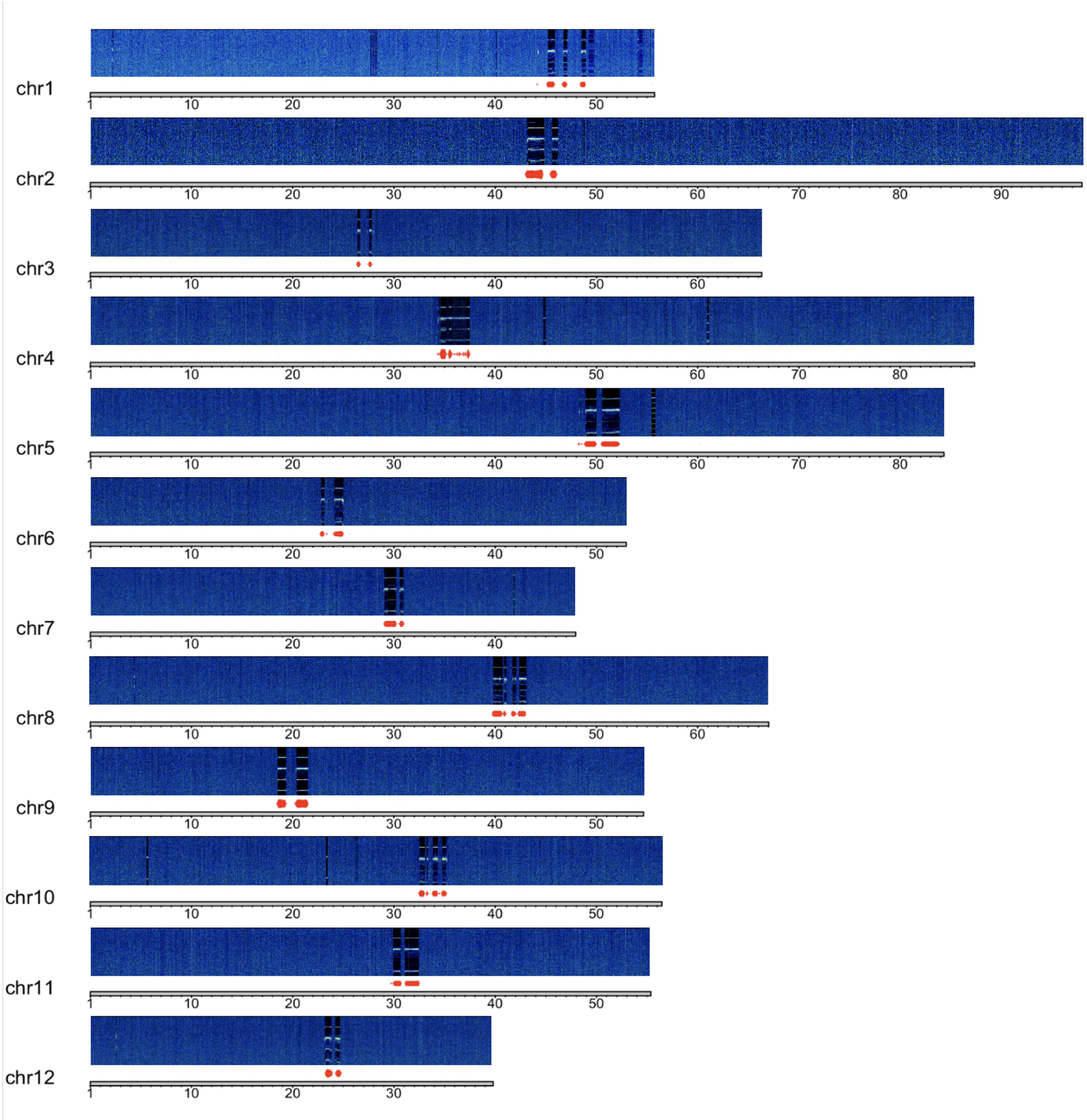


**Fig. S11. *Q. virginiana* centromeric calls.** The relative densities of the *QvCEN157* monomer (red) are plotted in combination with the chromosome-scale Fourier spectra generated by RepeatObserver. The x-axis of the spectra indicates genomic position and the y-axis of the spectra represents the frequency of a particular repeat and the corresponding color intensity showing the abundance of the repeat at that frequency in that position of the genome. The uppermost bar is described as the true or "fundamental" frequency and lower bars represent harmonics of that true frequency. The top bar across these species either correspond to 1/146 or 1/156:1/157, which agrees with the repeat monomers identified in *Q. virginiana* by TRASH. The presence of transposons can appear as "blurs" or "streaks" within and between the bars in the predicted array.

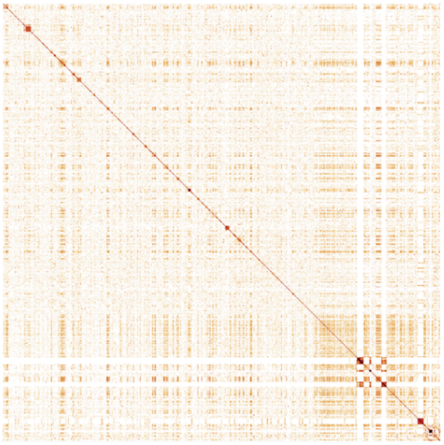

Chr 1 - Haplotype 1

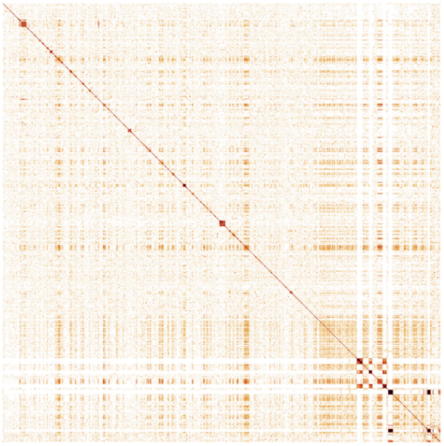

Chr 1 - Haplotype 2

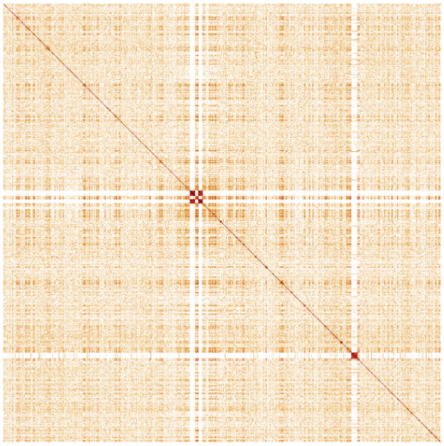

Chr 2 - Haplotype 1

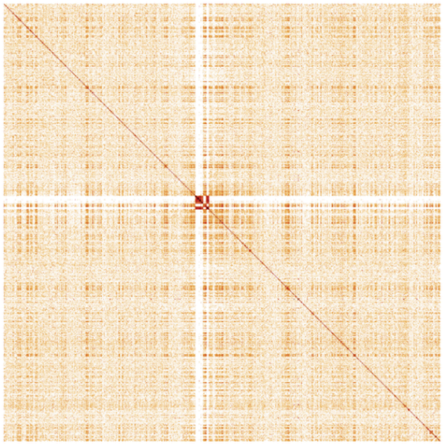

Chr 2 - Haplotype 2

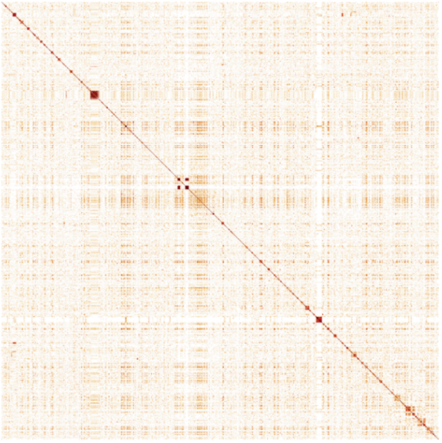

Chr 3 - Haplotype 1

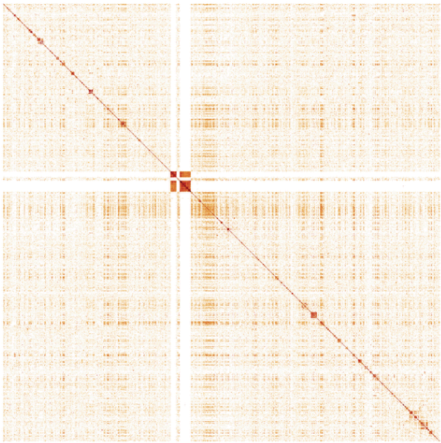

Chr 3 - Haplotype 2

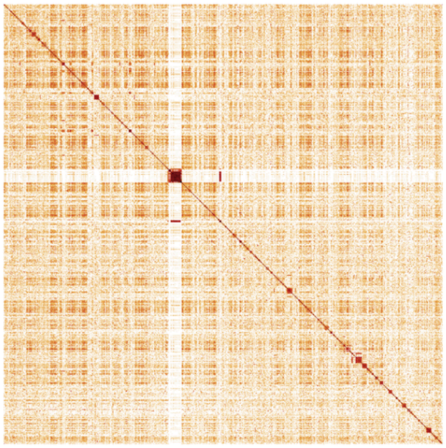

Chr 4 - Haplotype 1

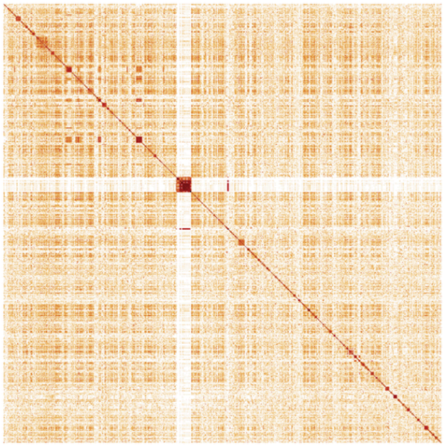

Chr 4 - Haplotype 2

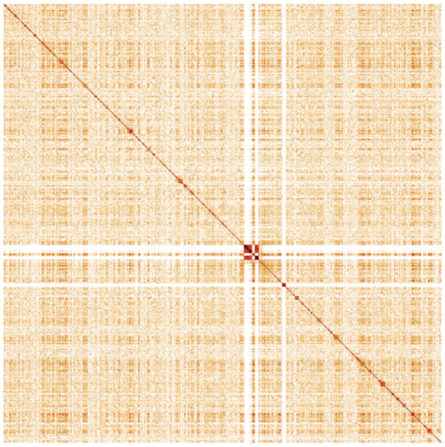

Chr 5 - Haplotype 1

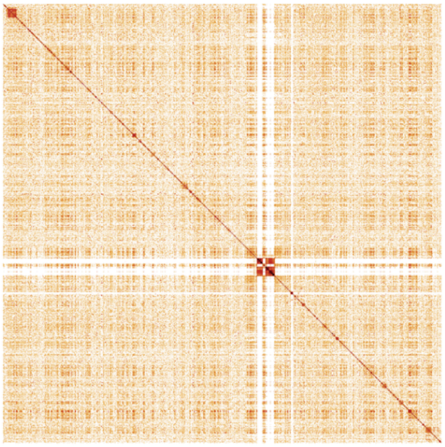

Chr 5 - Haplotype 2

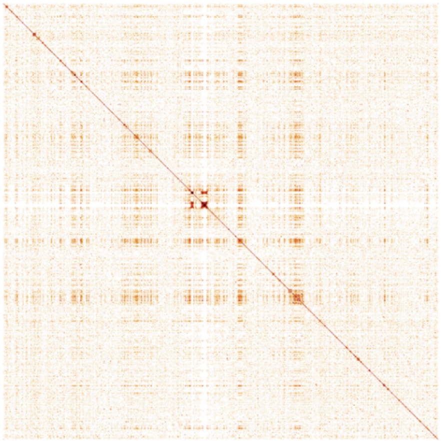

Chr 6 - Haplotype 1

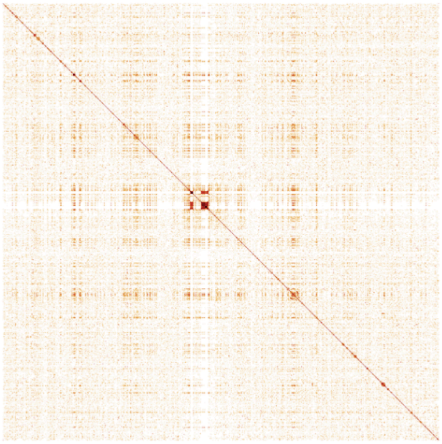

Chr 6 - Haplotype 2

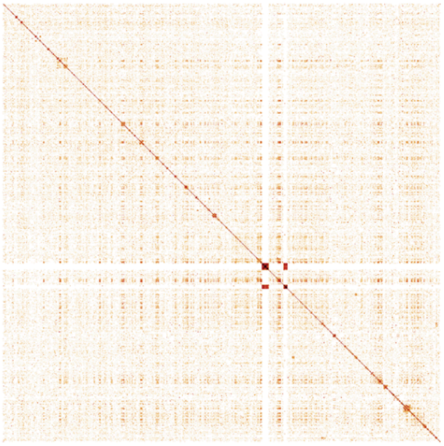

Chr 7 - Haplotype 1

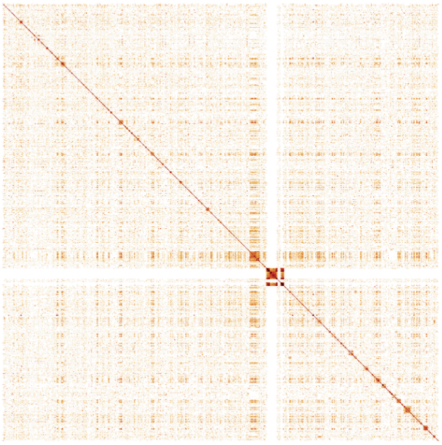

Chr 7 - Haplotype 2

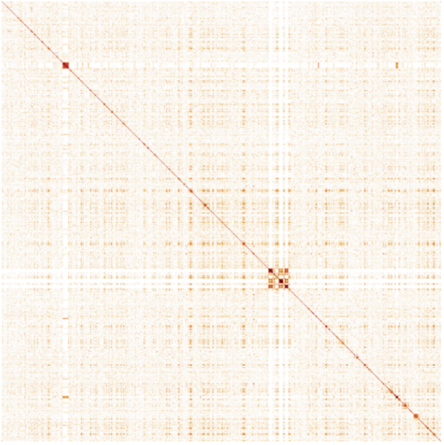

Chr 8 - Haplotype 1

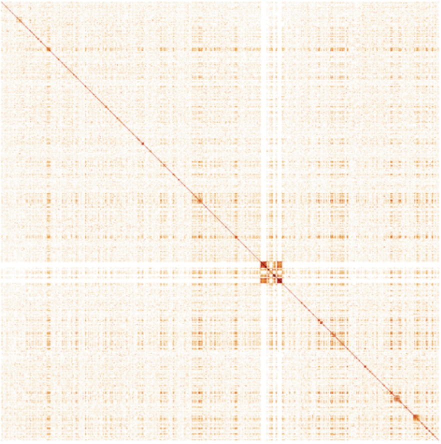

Chr 8 - Haplotype 2

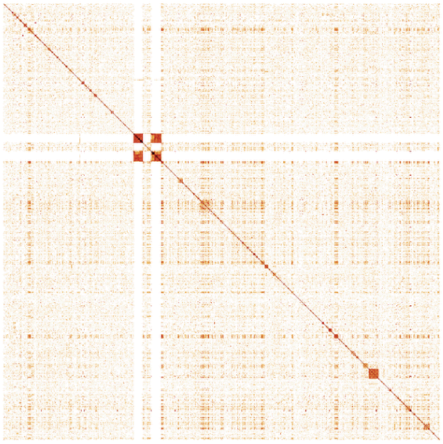

Chr 9 - Haplotype 1

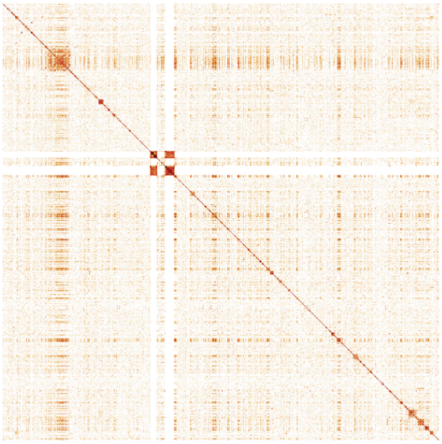

Chr 9 - Haplotype 2

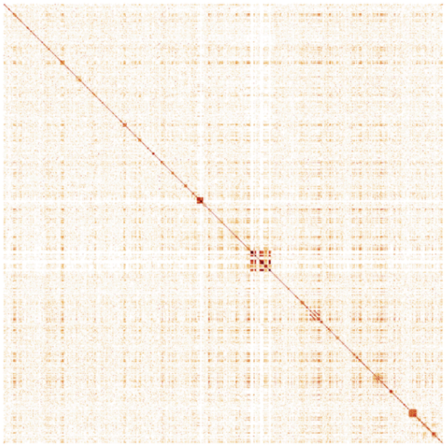

Chr 10 - Haplotype 1

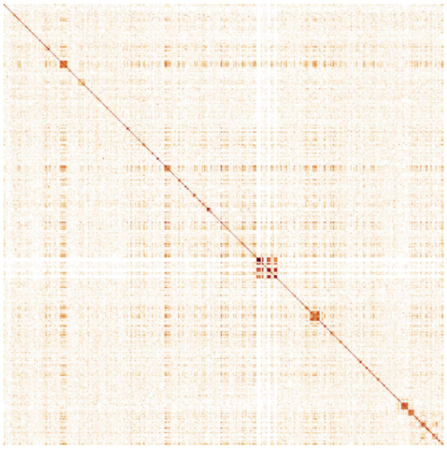

Chr 10 - Haplotype 2

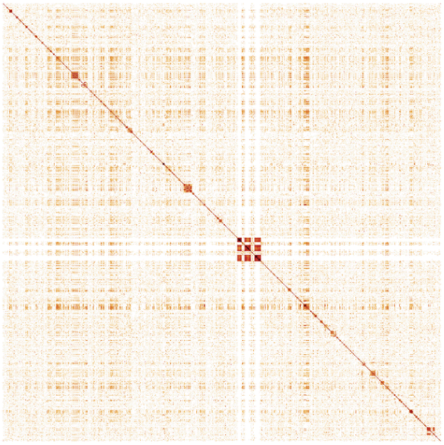

Chr 11 - Haplotype 1

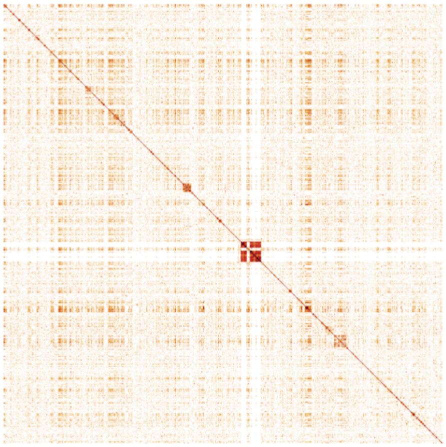

Chr 11 - Haplotype 2

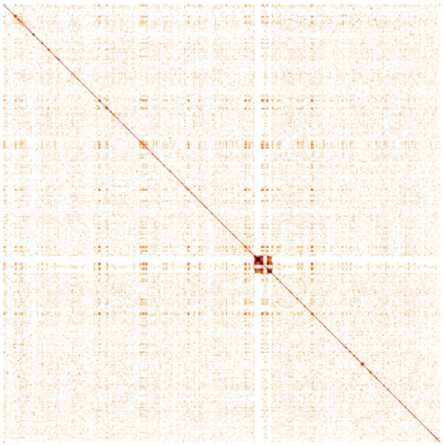

Chr 12 - Haplotype 1

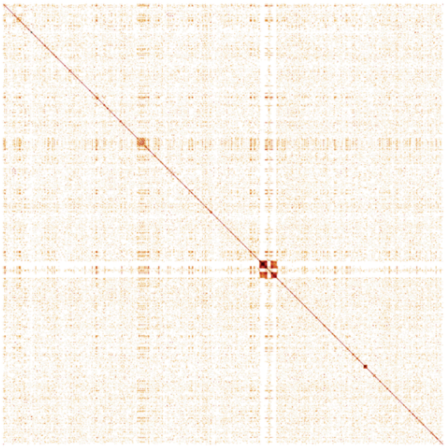

Chr 12 - Haplotype 2

**Fig. S12. StainedGlass identity heatmaps for visualizing the higher order structure and sequence relationships of repetitive elements in the chromosomes of both haplotypes.**

### *Q. alba* HapA

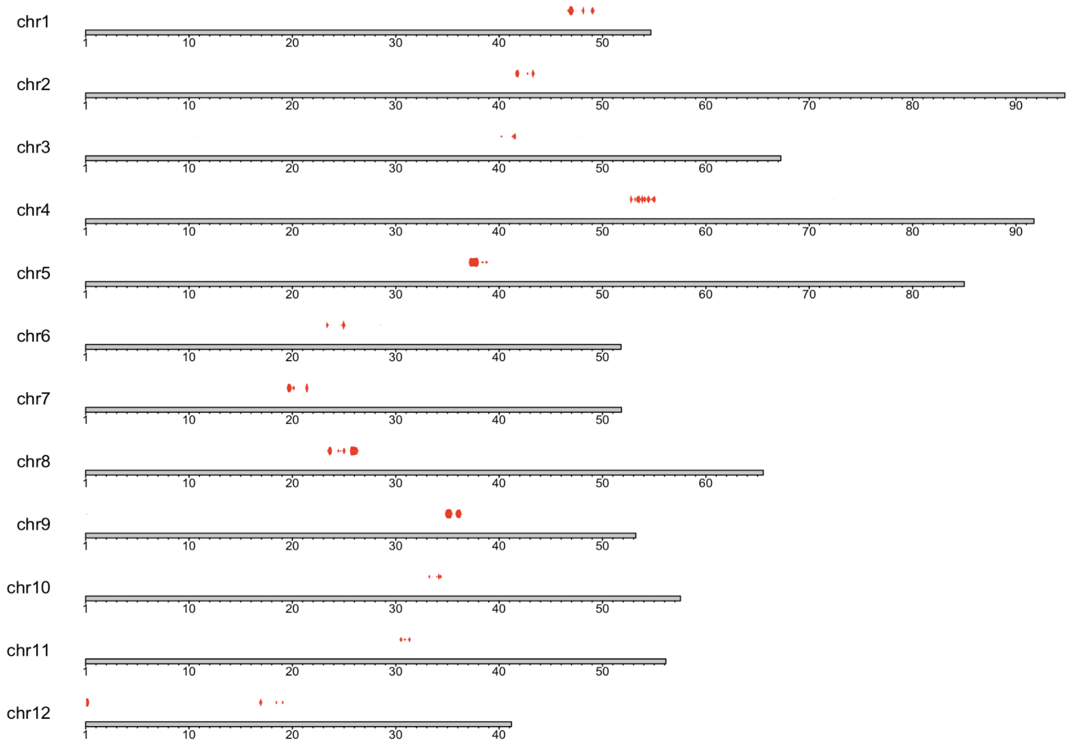


### *Quercus lobata*

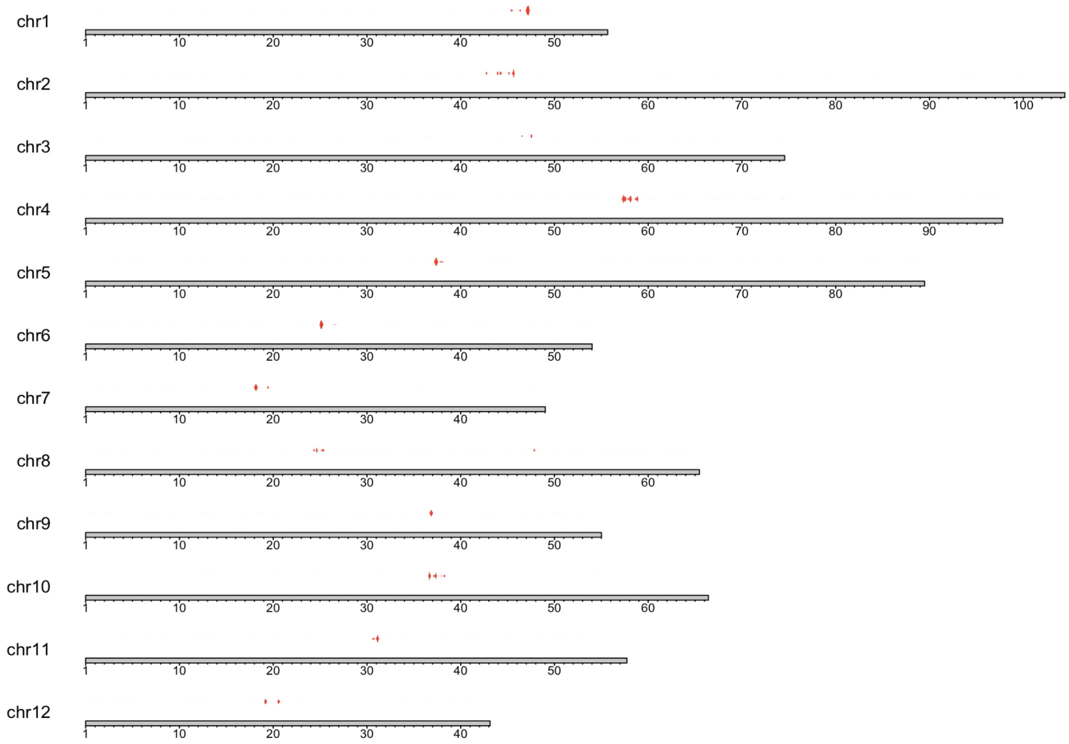

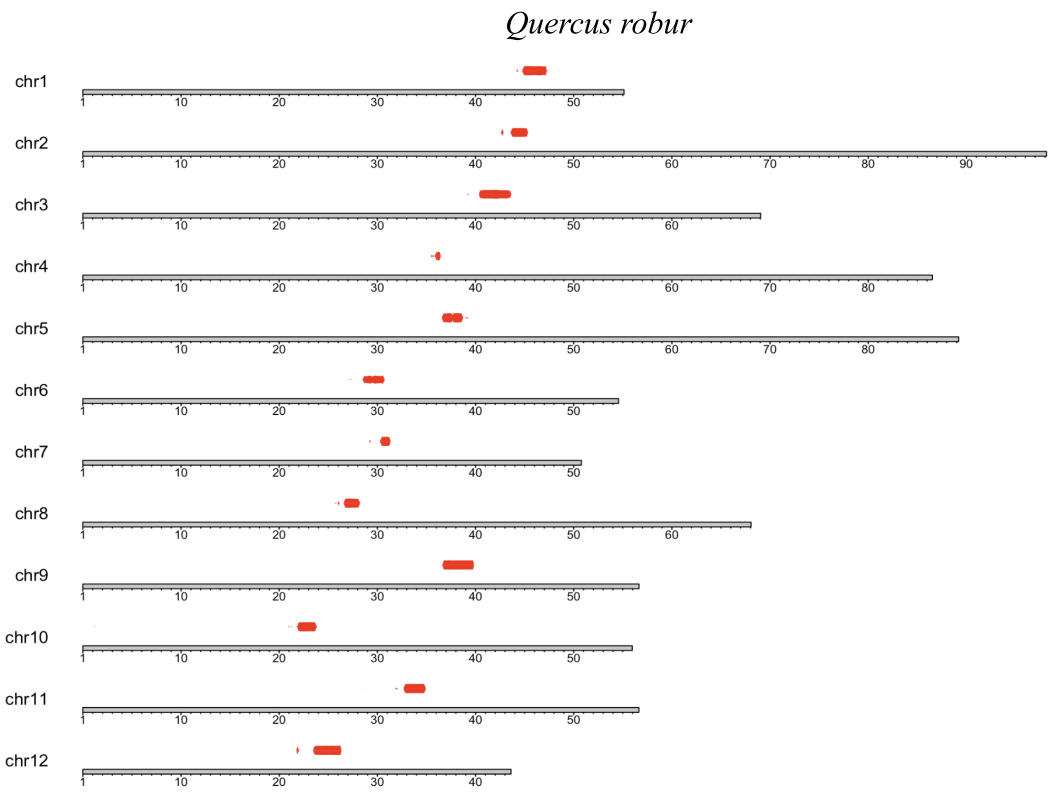


**Fig. S13, Blastn hits the *QvCEN157* monomer to other *Quercus* species.** A permissive blastn search with *-evalue 1*, *-word_size 4* and *-dust no* of *QvCEN157* against *Q. alba* HapA, *Q. lobata*, and *Q. robur* was performed. Although clearly indicating divergence relative to *Q. virginiana*, these alignments localize around the predicted centromeric locations for each species, with *Q. alba* and *Q. lobata* demonstrating a patchwork satellite pattern not shared by *Q. robur*.

```
qvirginiana_chr01_157       AAGTTGTTTTTTAATTTTTTGAATTTTTTTGCTATTTTTTTTCGGAATTT
qvirginiana_chr10_146_a     AAGTTGTTTTTTAATTTTTTGAATTTTTTTGCTATTTTTTT-CGGAATTT
                            ***************************************** ********

qvirginiana_chr01_157       GCTCCCCCGGGTCGAGTATGACGAGTATGAGCGGAATTCGGGTCTAAAAT
qvirginiana_chr10_146_a     TCTCCCCCGGGTCGAGTGT---------GAGCGGAATTCGGGTCTAAAAT
                             **************** *         **********************

qvirginiana_chr01_157       TCTTTTATCTCTTTCTCGGCCTATCTCATCCCCGTTTTGGCTAAAAATAA
qvirginiana_chr10_146_a     TCTTTTCTCTCTTTCTCGGCCTATCTCATCCT-GTTTTGGCTAAAAATAA
                            ****** ************************  *****************

qvirginiana_chr01_157       CGCCGGA
qvirginiana_chr10_146_a     TGCCGAA
                             **** *
```

**Fig. S14. MSA with CLUSTALW of *QvCEN157* and *QvCEN146* consensus sequences for Chromosome 1 of Haplotype 1**

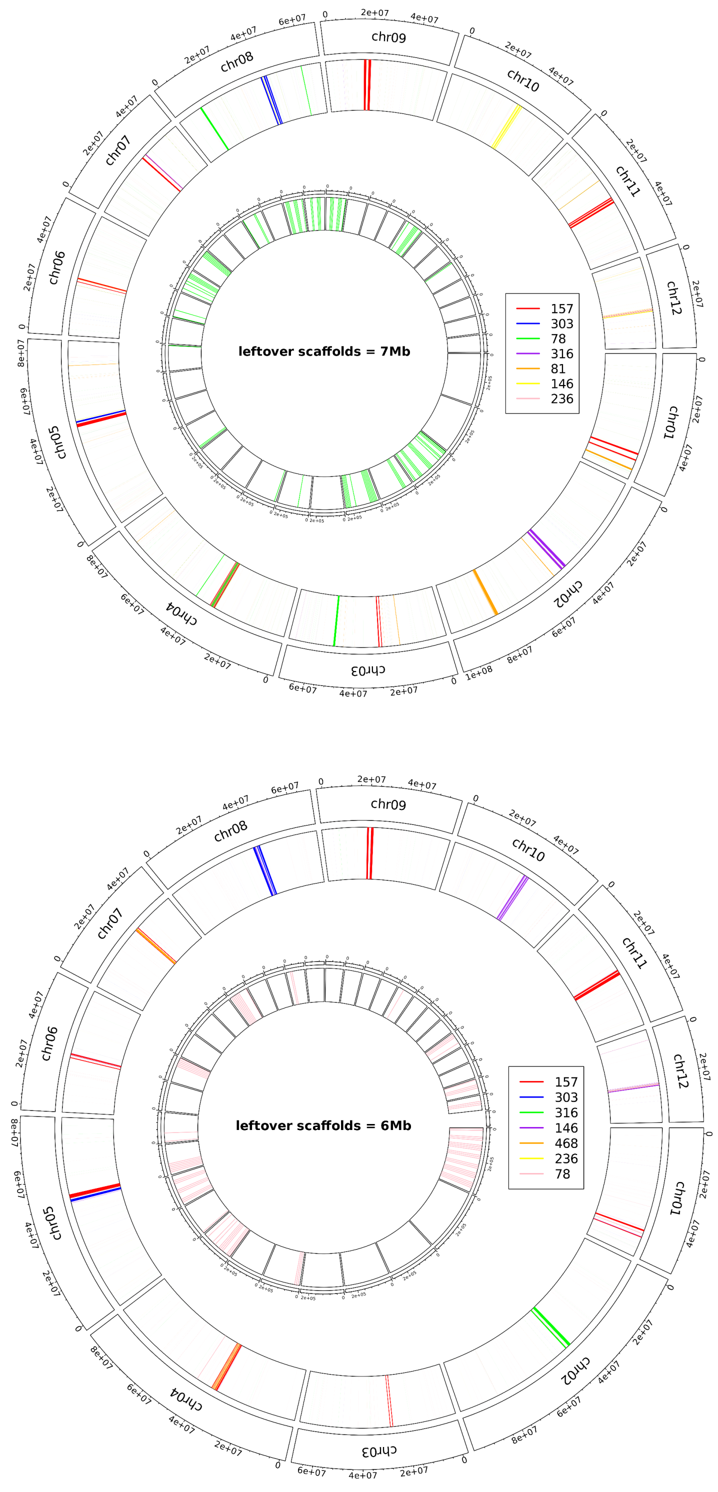


**Fig. S15. TRASH Circos plots for Haplotype 1 (top) and Haplotype 2 (bottom) of *Q. virginiana*.**

```
qvirginiana_chr01_157      AAGTTGTTTTTTAATTTTTTGAATTTTTTTGCTATTTTTTTTCGGAATTT
qvirginiana_chr10_146_a    AAGTTGTTTTTTAATTTTTTGAATTTTTTTGCTATTTTTTT-CGGAATTT
qlobata_CEN49              --------------------------------------GCTCATGGGCCC
                                                                   *   *

qvirginiana_chr01_157      GCTCCCCCGGGTCGAGTATGACGAGTATGAGCGGAATTCGGGTCTAAAAT
qvirginiana_chr10_146_a    TCTCCCCCGGGTCGAGTGTGA---------GCGGAATTCGGGTCTAAAAT
qlobata_CEN49              CCGACCCGAGTTAGAAAATTCAAAAAATAAATGCAAAAAAATTCTAAAAA
                            *  ***  * * **   *             * **      *******

qvirginiana_chr01_157      TCTTTTATCTCTTTCTCGGCCT-ATCTCATCCCCGTTTTGGCTAAAAATA
qvirginiana_chr10_146_a    TCTTTTCTCTCTTTCTCGGCCT-ATCTCATCCT-GTTTTGGCTAAAAATA
qlobata_CEN49              TTAAAAAACATCATCCAGGCTTCATTTCAAGACGAAAACGGGTCAGAGAC
                           *       *    **   *** * ** ***          ** * * *

qvirginiana_chr01_157      ACGCCGGA----------------------------
qvirginiana_chr10_146_a    ATGCCGAA----------------------------
qlobata_CEN49              AGGCCGAAAAATAGAGAACAAAAATTTCATTCCTAA
                           * **** *
```

**Fig. S16. MSA with CLUSTALW of *Q. lobata putative CEN146* sequence, *QvCEN157* consensus sequences for *Q. virginiana*** **Chromosome 1, *QvCEN146* consensus sequences for *Q. virginiana*** **Chromosome 10**

### Haplotype 1 Centromeric Repetitive Elements

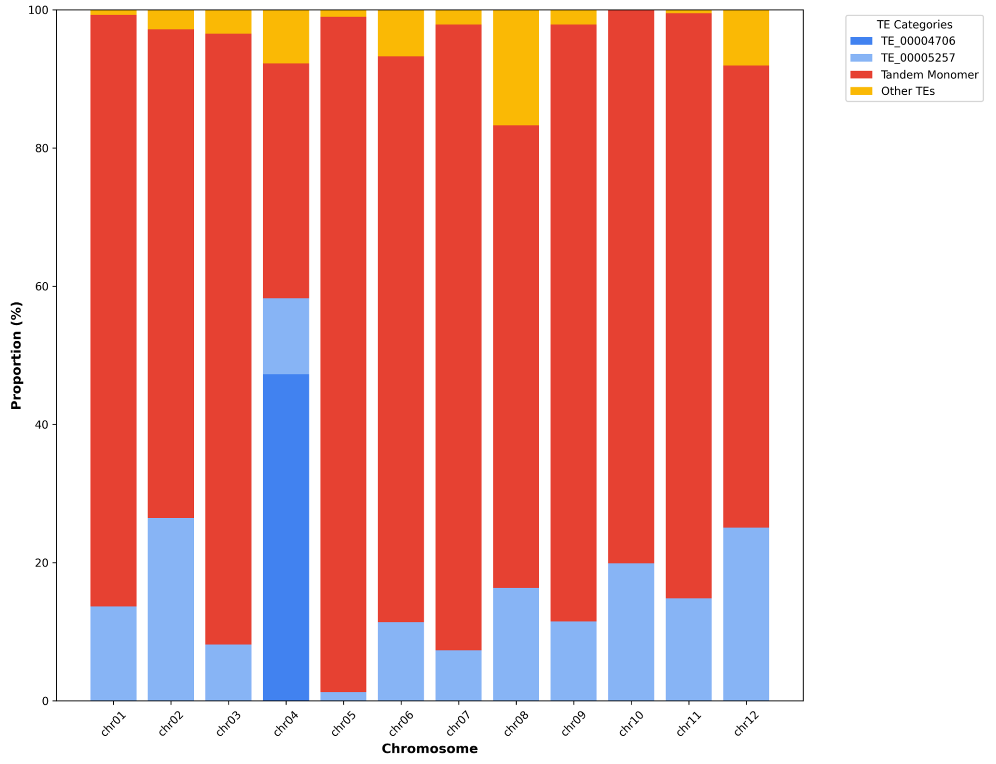


### Haplotype 2 Centromeric Repetitive Elements

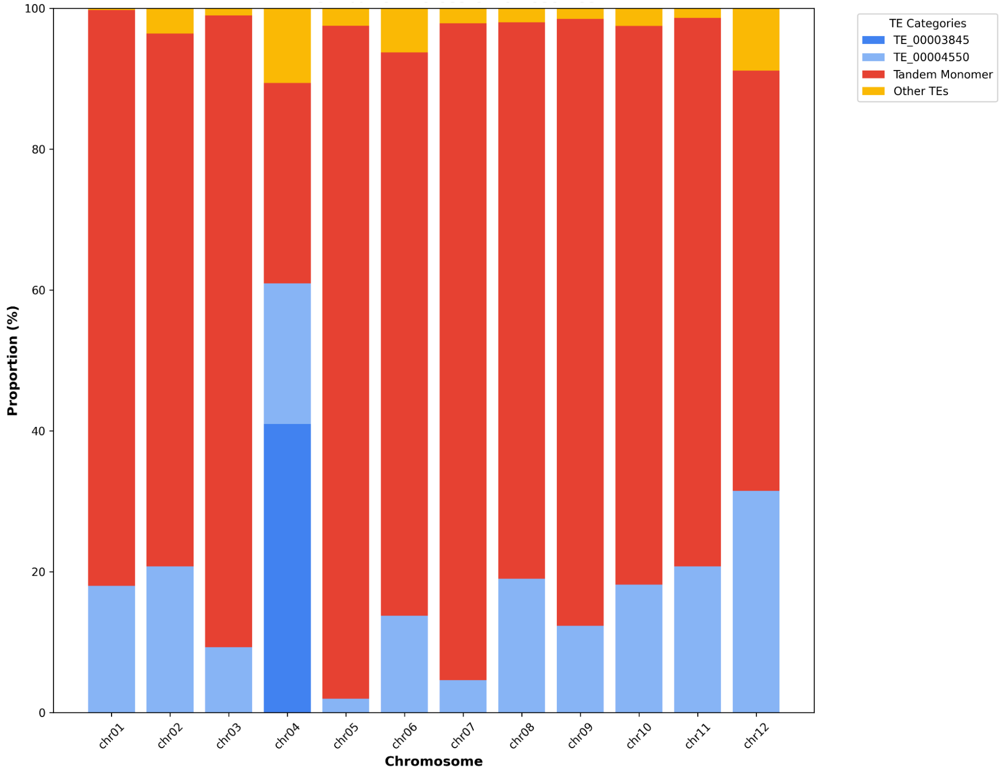

**Fig. S17. The proportion of the TEs identified by EDTA in the centromeric satellites of each chromosome.** The annotations for each of the ranges specified in Supplementary Table S7 were merged for each chromosome, then parsed and summarized using a custom python script. Although annotated as "Unknown" by EDTA, the TEs marked as tandem monomers in this figure were the most abundant annotations in these ranges (Hap1: TE_00000218, TE_00001084, TE_00000000; Hap2: TE_00000223) and were within the range of a typical centromeric monomer. Although similar between the haplotypes, they did not align as readily with the centromeric monomers identified by TRASH. This is likely due to differences in consensus sequence generation between the two programs.

### Supplemental Methods

#### S1. Identification, Characterization, and Visualization of the *Q. virginiana* Centromeres

Prior to trying to identify centromeres computationally, it can be helpful to determine if any previous karyotypic work has been conducted on your species of interest. In the case of *Q. virginiana*, no prior karyotypes were readily available. However, some centromere characterization had been performed on the haploid reference for *Q. lobata*, which we used to inform and anchor our own observations (Sork et al. 2022).

First, we generated interactive identity heatmaps for each chromosome with StainedGlass (Vollger et al. 2022 Jan 10). ModDotPlot can be used as a faster, but lower resolution alternative if needed (Sweeten et al. 2024). From these visualizations, we identified putative centromeric regions by visually scanning for areas containing high-complexity arrays. In StainedGlass plots, these regions typically appear as darker, more compact sections with high self-identity scores relative to other parts of the chromosome. It is important to note that these arrays do not exclusively represent centromeres, as other repeat-rich genomic sections can produce similar patterns. However, in genomes with high transposable element (TE) content in their centromeres and/or substantial tandem monomer content, this pattern tends to be characteristic. This feature is generally most visible in recently assembled genomes, as newer sequencing methods and assembly algorithms have improved the resolution and contiguity of centromeres. We have noticed that for genomes generated prior to the emergence of CCS HiFi sequencing or more unusual chromosomes such as sex chromosomes, these features are more difficult to detect.

Using the interactive HiGlass tool, we determined approximate boundaries of the putative centromeres to 10 kb resolution. Because biological boundaries of centromeres are defined by CENH3 localization, these computational "boundaries" represent estimates only. We combined this visualization-based approach with additional checks to further characterize these putative centromeric regions:

- **RepeatObserver** (Elphinstone et al. 2025): RepeatObserver is a powerful analytical tool that enables visualization of 3-5000bp repeats using a Fourier transform of DNA walks. The spectra that it generates can be used in combination with TRASH (explained below).
- **TRASH** (Wlodzimierz et al. 2023): TRASH is a novel method for identifying tandem repeats and their higher-order structures in diverse genomes. TRASH is particularly effective for identifying conserved centromeric monomers. This program generates multiple files useful for downstream analysis. We specifically utilized the "Summary.of.repetitive.regions_assembly.fa.csv" file, which provides information about monomer consensus sequences, and the "assembly_circos.pdf" file, which plots repeat monomers and their frequencies along chromosomes and scaffolds. The CSV file can be sorted to identify the most frequently occurring consensus monomers across chromosomes. Highly frequent tandem monomers falling within the 125-300 bp range represent excellent candidates for centromeric monomers, as this constitutes the current

standard range for most plant centromeric monomers (Melters et al. 2013). However, it is important to note that not all centromeres are characterized exclusively by tandem monomers and may also be invaded by (or entirely composed of) transposable elements (Naish and Henderson 2024). The monomer sizes identified by TRASH can be compared with the frequencies represented in the spectra generated by RepeatObserver.

- **TRF** (Benson 1999): Furthermore, we used TRF (Tandem Repeat Finder) to identify monomers greater than 100 bp, which has classically been the approach for identifying putative centromeres in *de novo* genome assemblies. As a quality check, we subset the output file by monomers with lengths greater than 100 bp, then plotted these to observe their density relative to putative centromeric regions and assess alignment.
- **NeSSie** (Berselli et al. 2018)**:** Centromeric regions typically exhibit distinct sequence symmetries relative to other regions on the chromosome. To visualize these characteristics, we calculated the Shannon entropy and linguistic complexity of each chromosome sequence using NeSSie. These calculations were performed using a sliding window approach (1 kb window lengths with 100 bp shifts). Entropy in centromeric regions (and other densely repetitive regions) is typically less variable than in other genomic regions, resulting in visible constrictions in plots at these locations. Similarly, linguistic complexity naturally decreases in highly repetitive regions, producing values below the genome average in these areas.

In addition to this, we also subset the TEanno.gff3 output from EDTA into the main superfamilies identified and plotted their relative densities to each chromosome to observe larger patterns surrounding or related to our regions of interest. We recommend plotting the density of different TE superfamilies with gene density along each chromosome as this approach can provide insights into broader genome architecture features, such as whether gene density decreases or certain repeats increase near suspected centromeric regions.

When additional genomes are available for your target species or related species within the genus, we recommend also performing this analysis on those genomes as a validation check to determine whether similar patterns and positioning for putative centromeres can be identified in other individuals. Karyotypes are often conserved between closely related species, and centromeric monomers may be sufficiently conserved to permit permissive BLAST searches between species without significant issues, as observed in many *Quercus* species. However, this conservation is highly species- and clade-specific.

We note that this approach has been applied exclusively to monocentric angiosperms and has not been tested on other species or clades, particularly those with holocentromeres or neocentromeres, where patterns may differ substantially.

**References –**

Benson G. 1999. Tandem repeats finder: a program to analyze DNA sequences. Nucleic Acids Res. 27(2):573–580.

Berselli M, Lavezzo E, Toppo S. 2018. NeSSie: a tool for the identification of approximate DNA sequence symmetries. Bioinformatics. 34(14):2503–2505.

Elphinstone C, Elphinstone R, Todesco M, Rieseberg LH. 2025 Mar 4. RepeatOBserver: Tandem repeat visualisation and putative centromere detection. Mol Ecol Resour.:e14084.

Melters DP, Bradnam KR, Young HA, Telis N, May MR, Ruby JG, Sebra R, Peluso P, Eid J, Rank D, et al. 2013. Comparative analysis of tandem repeats from hundreds of species reveals unique insights into centromere evolution. Genome Biol. 14(1):R10.

Naish M, Henderson IR. 2024. The structure, function, and evolution of plant centromeres. Genome Res. 34(2):161–178.

Sork VL, Cokus SJ, Fitz-Gibbon ST, Zimin AV, Puiu D, Garcia JA, Gugger PF, Henriquez CL, Zhen Y, Lohmueller KE, et al. 2022. High-quality genome and methylomes illustrate features underlying evolutionary success of oaks. Nat Commun. 13(1):2047.

Sweeten AP, Schatz MC, Phillippy AM. 2024. ModDotPlot-rapid and interactive visualization of tandem repeats. Bioinformatics. 40(8). doi:10.1093/bioinformatics/btae493. http://dx.doi.org/10.1093/bioinformatics/btae493.

Vollger MR, Kerpedjiev P, Phillippy AM, Eichler EE. 2022 Jan 10. StainedGlass: Interactive visualization of massive tandem repeat structures with identity heatmaps. Bioinformatics. doi:10.1093/bioinformatics/btac018. http://dx.doi.org/10.1093/bioinformatics/btac018.

Wlodzimierz P, Hong M, Henderson IR. 2023. TRASH: Tandem Repeat Annotation and Structural Hierarchy. Bioinformatics. 39(5):btad308.

### S2. Investigations into HGT in *Q. virginiana*

To investigate potential horizontal gene transfer (HGT) into *Q. virginiana*, we searched for lateral transfers from the cynipid gall wasp *Belonocnema kinseyi*, a gall-inducing wasp hosted by *Q. virginiana*, as well as bacterial and fungal sources. In each analysis, protein sequences from *Q. virginiana* were compared with those from either *B. kinseyi*, NCBI RefSeq bacterial proteins, or NCBI RefSeq fungal proteins (sourced on September 12, 2024), as well as *Arabidopsis thaliana* (GCF_000001735.4/TAIR10.1) and *Populus trichocarpa* (GCF_000002775.5/P.trichocarpa_v4.1). A reciprocal analysis was conducted on the genome of *B. kinseyi* to detect lateral transfers from oak to wasp. This analysis used *Q virginiana* as the potential donor and compared similarity to *Apis mellifera* (GCF_003254395.2/Amel_HAv3.1) and *Nasonia vitripennis* (GCF_009193385.2/Nvit_psr_1.1).

BLASTP searches were conducted to identify oak or wasp proteins with higher similarity to potential donor lineages than to other plants or wasps, using the following filtering criteria. Sequences were considered candidates if they shared more than 75% of their aligned amino acid residues, to exclude coincidental resemblances; were less than 50% identical to proteins in close relatives and at least 20% more identical to a donor sequence than to any relative sequence, to exclude conserved lineage-specific proteins; and were no more than 90% identical overall, to exclude universal housekeeping genes shared across eukaryotes.

For the *Q. virginiana* and *B. kinseyi* genomes, an additional analysis was conducted at the nucleotide level to discover transfers of sequences not included in the protein annotations of each genome. Using KMC3 (v3.2.4), all possible 31-nucleotide k-mers from both genomes were generated and compared to identify exact matches between species. Shared k-mers were then mapped back to their source genomes using BWA-mem to identify regions with clusters of matches. Clusters were defined as regions containing at least 3 k-mers within 50 base pairs of each other, spanning at least 100bp, with a minimum density of 0.1 k-mers per base pair. To exclude potential false positives, sequences from identified clusters were extracted and compared against the NCBI nucleotide database.

Horizontal gene transfer into plants is so far known only from *Agrobacterium tumefaciens*, a long-term endosymbiotic parasite known to manipulate its host's genome. *Q. virginiana* is a long-term host of many endosymbiotic bacteria and fungi, some of which, like *Taphrina*, are known to induce morphological changes in plant tissue. Gall-inducing wasps like *Belonocnema* create tissues made through the differential expression of plant genes and therefore possess mechanisms to interface directly or indirectly with the host genome. Transfer from host plants into specialist herbivorous insects has been documented in at least 14 species, including one Hymenopteran (Li et al 2022), and may facilitate the evolution of more effective herbivory (Gilbert and Maumus, 2023). Since the exact mechanisms by which gall wasps affect gene regulation in oaks remain unclear, it cannot be ruled out that such mechanisms may have provided opportunities for gene transfer during the evolution of the symbiosis.

No strong candidates for HGT were identified in the *Q. virginiana* or *B. kinseyi* genomes. BLASTP comparisons of oak proteins against *B. kinseyi* yielded no proteins with significantly higher similarity to the wasp genome than to plant references, and vice versa. Similarly, analyses of oak proteins against bacterial sequences from NCBI RefSeq revealed no candidates meeting the filtering criteria. One oak protein, mitochondrial protoheme IX farnesyltransferase, exhibited higher similarity to a fungal sequence than to plant references, but this result is likely a false positive, as the protein is a conserved mitochondrial enzyme without clear biological context for HGT. A k-mer based analysis of the complete *Q. virginiana* and *B. kinseyi* genomes identified multiple regions containing exact 31bp matches between oak and wasp, but manual inspection revealed these included only simple sequence repeats and one conserved large subunit ribosomal RNA sequence. No unique gene sequences or potential horizontally transferred regions were identified using this approach, suggesting that the absence of detected HGT between wasp and oak is not due to limitations in gene prediction or annotation. These findings suggest that, if HGT into the *Q. virginiana* genome from these sources has occurred, it involves sequences not captured by this filtering approach.

**References–**

Li Y, Liu Z, Liu C, Shi Z, Pang L, Chen C, Chen Y, Pan R, Zhou W, Chen XX, Rokas A. 2022. HGT is widespread in insects and contributes to male courtship in lepidopterans. Cell. 185(16):2975-2987. Available from: https://www.sciencedirect.com/science/article/pii/S009286742200719X?via%3Dihub#mmc2

Gilbert C, Maumus F. 2023. Sidestepping Darwin: horizontal gene transfer from plants to insects. Curr Opin Insect Sci. 57:101035. Available from: https://www.sciencedirect.com/science/article/pii/S2214574523000329
